# Functions for Cdc42p BEM Adaptors in Regulating a Differentiation-Type MAP Kinase Pathway

**DOI:** 10.1101/786483

**Authors:** Sukanya Basu, Beatriz González, Boyang Li, Garrett Kimble, Keith G. Kozminski, Paul J. Cullen

**Author notes:** corresponding author Address: 532 Cooke Hall, Department of Biological Sciences, State University of New York at Buffalo, Buffalo, NY 14260-1300. Author contributions: S.B. designed experiments, generated data, and wrote the paper; B.G. performed experiments and analyzed the data; B.L. generated data; G.K. generated data; K.K. provided reagents, generated data, and analyzed the data; P.J.C. designed experiments and wrote the paper. The authors have no competing interests in the study. **Impact Sentence:** Rho GTPase adaptors play unique and sequential roles in the regulation of MAPK pathways.

## Abstract

Rho GTPases regulate cell polarity and signal transduction pathways to control morphogenetic responses in different settings. In yeast, the Rho GTPase Cdc42p regulates cell polarity, and through the p21-activated kinase Ste20p, Cdc42p also regulates mitogen-activated protein kinase (MAPK) pathways (mating, filamentous growth or fMAPK, and HOG). Although much is known about how Cdc42p regulates cell polarity and the mating pathway, how Cdc42p regulates the fMAPK pathway is not clear. To address this question, Cdc42p-dependent MAPK pathways were compared in the filamentous (∑1278b) strain background. Each MAPK pathway showed a unique activation profile, with the fMAPK pathway exhibiting slow activation kinetics compared to the mating and HOG pathways. A previously characterized version of Cdc42p, Cdc42p^E100A^, that is specifically defective for fMAPK pathway signaling, was defective for interaction with Bem4p, the pathway-specific adaptor for the fMAPK pathway. Corresponding residues in Bem4p were identified that were required for interaction with Cdc42p and fMAPK pathway signaling. The polarity adaptor Bem1p also regulated the fMAPK pathway. In the fMAPK pathway, Bem1p recruited Ste20p to the plasma membrane, cycled between an open and closed conformation, and interacted with the GEF for Cdc42, Cdc24p. Bem1p also regulated effector pathways in different ways, behaving as a multi-functional adaptor in some pathways and an inert scaffold in others. Genetic suppression tests showed that Bem4p and Bem1p regulate the fMAPK pathway in an ordered sequence. Collectively, the study demonstrates unique and sequential functions for Rho GTPase adaptors in regulating MAPK pathways.

**HIGHLIGHTS:**

- Comparing Cdc42p-dependent MAPK pathways showed that the fMAPK pathway had slow activation kinetics compared to the mating and HOG pathways.
- A collection of *cdc42* alleles was tested for MAPK pathway functions.

§ Cdc42p^E100A^, previously characterized as being specifically defective for fMAPK signaling, showed reduced interaction with the fMAPK pathway adaptor Bem4p.
§ Corresponding residues in Bem4p were identified that were required for interaction with Cdc42p and fMAPK signaling.
- The polarity adaptor Bem1p regulated the fMAPK pathway.

§ Bem1p regulated the fMAPK pathway by recruiting Ste20p to the plasma membrane, cycling between an open and closed conformation, and interacting with the Cdc42p GEF, Cdc24p.
- Different domains of Bem1p had different roles in regulating effector pathways.

§ Bem1p may function as a multi-functional adaptor in some pathways and an inert scaffold in others.
- Bem4p and Bem1p regulated the fMAPK pathway in an ordered sequence.

§ The data support a model where Bem4p recruits Cdc24p to GDP-Cdc42p, and Bem1p directs GTP-Cdc42p to Ste20p at the plasma membrane.
§ The bud-site GTPase Rsr1p regulates Cdc24p in the fMAPK pathway but does not initiate signaling.

## INTRODUCTION

Mitogen-Activated Protein Kinase (MAPK) pathways are evolutionarily conserved signaling modules that control cell proliferation, cell differentiation, and the response to stress in eukaryotes (*1*). In the budding yeast *Saccharomyces cerevisiae*, MAPK pathways control cell differentiation to specific cell types, by the mating and filamentous growth or fMAPK pathways. MAPK pathways also control the response osmotic stress, by the high osmolarity glycerol response (HOG) pathway, and the response to compromised cell integrity by the protein kinase C pathway. Like many signaling pathways, yeast MAPK pathways share components (*2–7*). For example, the Rho GTPase Cdc42p regulates three MAPK pathways (mating, fMAPK, and HOG). Cdc42p is also an essential protein that controls the establishment of cell polarity (*8*). How Cdc42p and other proteins induce pathway-specific outputs in a functionally interconnected network is not clear. This question is relevant because in higher organisms, cross-talk and mis-regulation of GTPase pathways, like those requiring CDC42 and RAS, is a leading cause of cancer and other diseases. Understanding how common proteins function in pathway-specific settings remains at the forefront of studies of signaling pathway regulation (*9–12*).

In response to carbon or nitrogen limitation, yeast and other fungal species can undergo a differentiation response called filamentous or invasive/pseudohyphal growth (*13, 14*). In some plant and animal pathogens, like the major human pathogen *Candida albicans*, filamentous growth is required for virulence (*15–18*). In yeast, filamentous growth occurs in wild strain backgrounds (Σ1278b is used here) because the properties that are associated with filamentous growth have been lost due to genetic manipulation of laboratory strains (*19–22*). Although hundreds of genes contribute to filamentous growth, the response is mainly characterized by the formation of elongated cells that grow in distal-unipolar budding pattern and remain attached to each other to form chains. The Cdc42p-dependent fMAPK pathway is one of the major signaling pathways that regulates filamentous growth (ROBERTS AND FINK 1994; CULLEN *et al.* 2004; CULLEN AND SPRAGUE 2012). Studies of the fMAPK pathway have provided insights into how ERK-type MAPK pathways regulate eukaryotic cell differentiation and fungal pathogenesis.

The fMAPK pathway shares components with the HOG and mating pathways, yet each pathway induces a specific set of target genes and a unique response (*3-5, 13, 23-27*). A core module composed of Cdc42p, p21-activated kinase (PAK) Ste20p (*23, 24*), and MAPKKK Ste11p regulate all three MAPK pathways. This core module regulates different MAP kinases for the three pathways. The MAP kinase Kss1p is the main regulator of the fMAPK pathway (*28*). In the fMAPK pathway, Kss1p regulates a suite of transcriptional activators (Ste12p, Tec1p, Msa1p and Msa2p) and repressors (Dig1p and Dig2p) (*28–35*) to activate transcription at filamentation-specific promoters (*34, 36*). Tec1p in particular functions exclusively in the fMAPK pathway and is degraded during the mating response (*37, 38*).

Pathway-specific factors that function at the plasma membrane also regulate the Cdc42p module. The mucin-type glycoprotein Msb2p interacts with the GTP-bound conformation of Cdc42p to regulate the fMAPK pathway (*39*). Msb2p functions with two other transmembrane proteins, Sho1p and Opy2p (*40–49*). Msb2p, Sho1p, and Opy2p regulate the fMAPK pathway and also the HOG pathway but not the mating pathway.

In addition to Msb2p, Bem4p is a Cdc42p-interacting protein (*50, 51*) that regulates the fMAPK pathway. Bem4p regulates the fMAPK pathway but not the HOG or mating pathways (*52*). Bem4p associates with Cdc42p, the guanine nucleotide exchange factor (GEF) Cdc24p (*53–55*), the MAPKKK Ste11p, and the MAPK Kss1p (*52*). Thus, Bem4p might direct Cdc42p, in some manner, to a pathway-specific complex containing Kss1p. Finally, bud-site-selection proteins, culminating on the Ras-type GTPase Rsr1p (*56, 57*) also regulate the fMAPK pathway but no other Cdc42p-dependent MAPK pathways (*58*). Therefore, at least three Cdc42p-interacting proteins, Msb2p, Bem4p, and Rsr1p regulate the fMAPK pathway.

Despite the utility of studying MAPK pathways in a genetically amenable system, Cdc42p-dependent MAPK pathways have not been directly compared in yeast. One reason is that most studies of the mating and HOG pathways have been carried out in laboratory strains, which have lost the ability to undergo filamentous growth (*19, 21, 22*). Another reason is that the HOG pathway has redundant branches that converge on the MAPKK Pbs2p, bypassing Cdc42p. Only one branch is dependent on Cdc42p, the Ste11p branch (*59–61*). By directly comparing Cdc42p-dependent MAPK pathways in yeast, we show that the pathways have different kinetics. We also found that the interaction between Bem4p and Cdc42p is critical for fMAPK pathway signaling. We also show that the major polarity adaptor for Cdc42p, Bem1p, also regulates the fMAPK pathway. Bem1p also regulates the mating (*62*) and HOG pathways (*63*), so its role in regulating the fMAPK pathway might be expected. What was surprising was that Bem1p regulated different effector pathways in different ways. Bem1p regulated the fMAPK pathway by recruiting Ste20p to the plasma membrane, cycling between an open and closed conformation, and interacting with Cdc42p’s GEF, Cdc24p. We propose a model for how the Cdc42p module is directed to the fMAPK pathway, by interactions with Cdc24p (by Bem4p, Bem1p and Rsr1p), Cdc42p (by Bem4p and Bem1p), and Ste20p (by Bem1p). Our study provides a specific mechanism for how Cdc42p is activated in the context of a differentiation-type MAPK pathway.

## RESULTS

### Signaling Landscape of Cdc42p-Dependent MAPK Pathways

Three MAPK pathways in yeast (fMAPK, HOG, and mating) require common components yet induce different responses [Fig. 1A, (*27, 64, 65*)]. The Rho-type GTPase Cdc42p, PAK Ste20p, and MAPKKK Ste11p are central to the regulation of the three pathways (Fig. 1A, grey box). To directly compare Cdc42p-dependent MAPK pathways, experiments were performed in filamentous (∑1278b) strains lacking the Sln1p-branch of the HOG pathway (*ssk1*Δ). Growth of cells in the non-preferred carbon source galactose (GAL) induced filamentous growth (Fig. 1B, left), phosphorylation of the MAP kinase Kss1p (Fig. 1B, right, P∼Kss1p and to a minor extent P∼Fus3p), and expression of the p*FRE-lacZ* reporter (Fig. 1E, red). Osmotic shock (0.4 M KCl) did not induce a morphogenetic response (Fig. 1C, left), but induced rapid phosphorylation of Hog1p (Fig. 1C, right) and expression of the p*8XCRE-lacZ* reporter (Fig. 1E, green). The mating pheromone α-factor induced mating projections (Fig. 1D, left), phosphorylation of Fus3p and Kss1p (Fig. 1D, right, note low levels of control protein at the 6 h time point), and expression of p*FUS1-lacZ* (Fig. 1E, blue). Kss1p is normally phosphorylated in response to α-factor and functions to modulate the mating response (*66, 67*).

**Figure 1.**
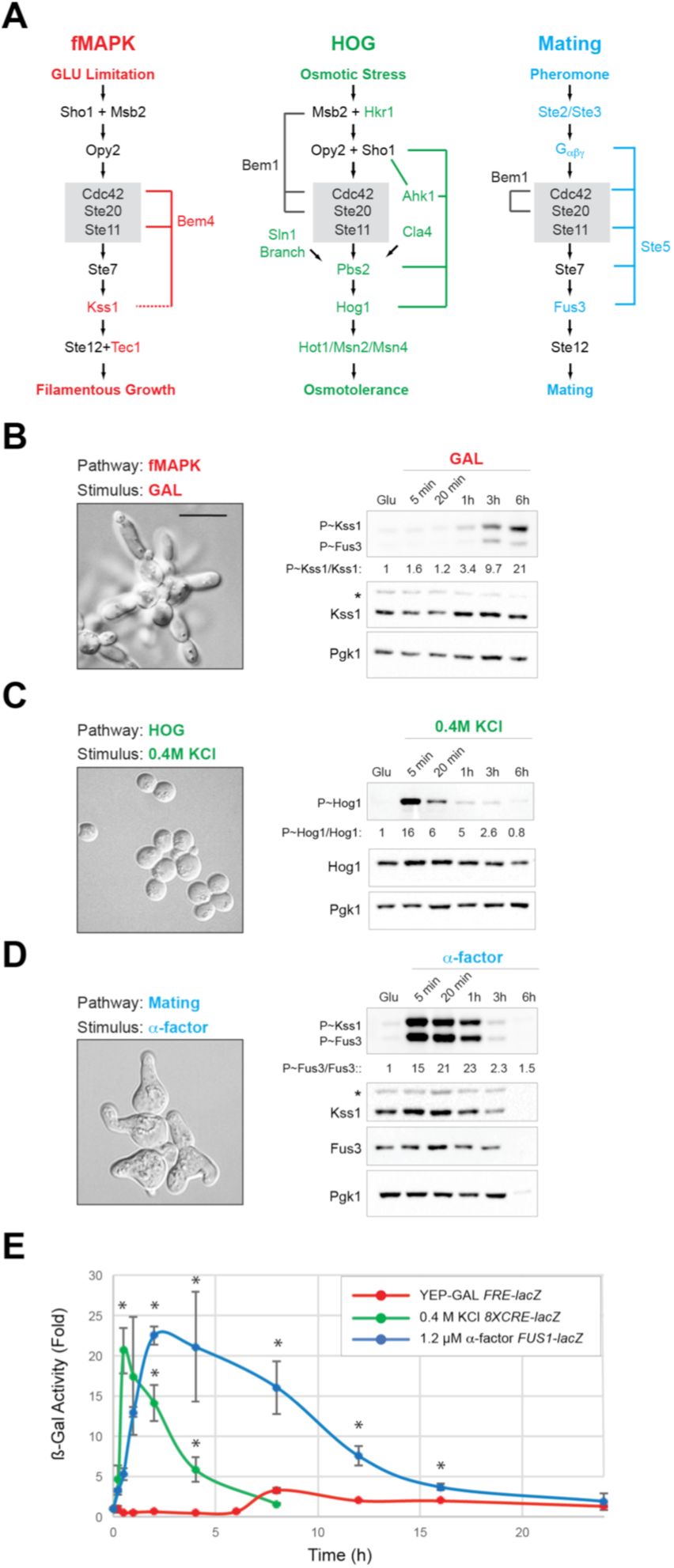
Comparison of MAP kinase pathways that require Cdc42p and other proteins. **A)** Three Cdc42p-dependent MAP kinase pathways. In color, pathway-specific factors, fMAPK (red), mating (blue), HOG (green); grey box, common or shared factors in all three pathways; black text, common to at least two pathways. **B)**. To assess fMAPK activity, wild-type cells (PC6810) were grown for 8h in YEP-GAL and photographed to generate the image at right (DIC 100X, bar, 10 microns). Immunoblots were probed with p44/p42 antibodies for P∼Kss1p and P∼Fus3p and with specific antibodies to detect Kss1p and Pgk1p. **C)** Wild-type cells were exposed to YPED + 0.4 M KCl (HOG) for 8h and photographed (Right, DIC 100X, bar, 10 microns). Antibodies to detect P∼Hog1p (p38), and Hog1p and Pgk1p as a control for proteins levels, were used. **D)** Wild-type cells were grown for 8h in YPED + 1.2 µM α-factor to induce mating response (Right, DIC 100X, bar, 10 microns). During the indicated times, samples were collected and probed with p44/42, Fus3p, Kss1p, and Pgk1p antibodies. In panels B, C and D, numbers indicate the relative band intensity of P∼MAPK to total MAPK levels normalized to the control, which was set to 1. Asterisk, a background band seen in some blots. **E)** Relative activity in fold difference compared to the uninduced condition (YEPD) for each reporter of the indicated transcriptional (*lacZ* fusion) reporters. Cells were grown as described in indicated media. β-galactosidase activity is shown for the times indicated. Each time point is the average of three independent trials. Error bars show the standard difference between experiments. Asterisk, p-value < 0.05 for values compared to *FRE-lacZ* for the times indicated.

The above experiments also showed that the fMAPK pathway was activated more slowly than the mating and HOG pathways. By P∼blot analysis, the fMAPK pathway was maximally activated by 6 h, compared to 5 min for HOG, and 1 h for the mating pathway (Fig. 1B-D). The mating and fMAPK pathways were directly comparable because P∼Fus3p and P∼Kss1p are recognized by the same antibodies (Fig. S1). By transcriptional reporter assays, the fMAPK pathway showed maximal activity by 7.5 h compared to 5 min for HOG and 20 min for mating pathways (Fig. 1E). The fMAPK pathway also showed less transcriptional reporter induction (4-fold for the *FRE-lacZ* reporter) than the HOG (20-fold) and mating pathways (20-fold). This observation is consistent with previous studies from our lab of the *FRE-lacZ* reporter (*68*) and other reporters that show moderate induction in GAL [e.g. *KSS1, YLRO42c, MSB2* (*49*)]. Therefore, the fMAPK pathway has slow activation kinetics compared to the HOG and mating pathways.

The activity of the Cdc42p-dependent MAPK pathways was also examined in response to exposure to their non-cognate stimuli (Figs. S1-2). With two exceptions, the pathways functioned in an insulated manner: 1) GAL induced the HOG pathway at a rate far less than that observed with KCl induction (Fig. S1A, p38 blot), which occurs during the filamentation response (*69*); and 2) α-factor induced phosphorylation of Hog1p (Fig. S1A, p38 blot), which did not result in induction of the *8XCRE–lacZ* reporter (Fig. S2B). These results support the prevailing view that Cdc42p-dependent pathways function in an insulated manner to induce specific transcriptional and morphogenetic responses to specific stimuli.

**Figure 2.**
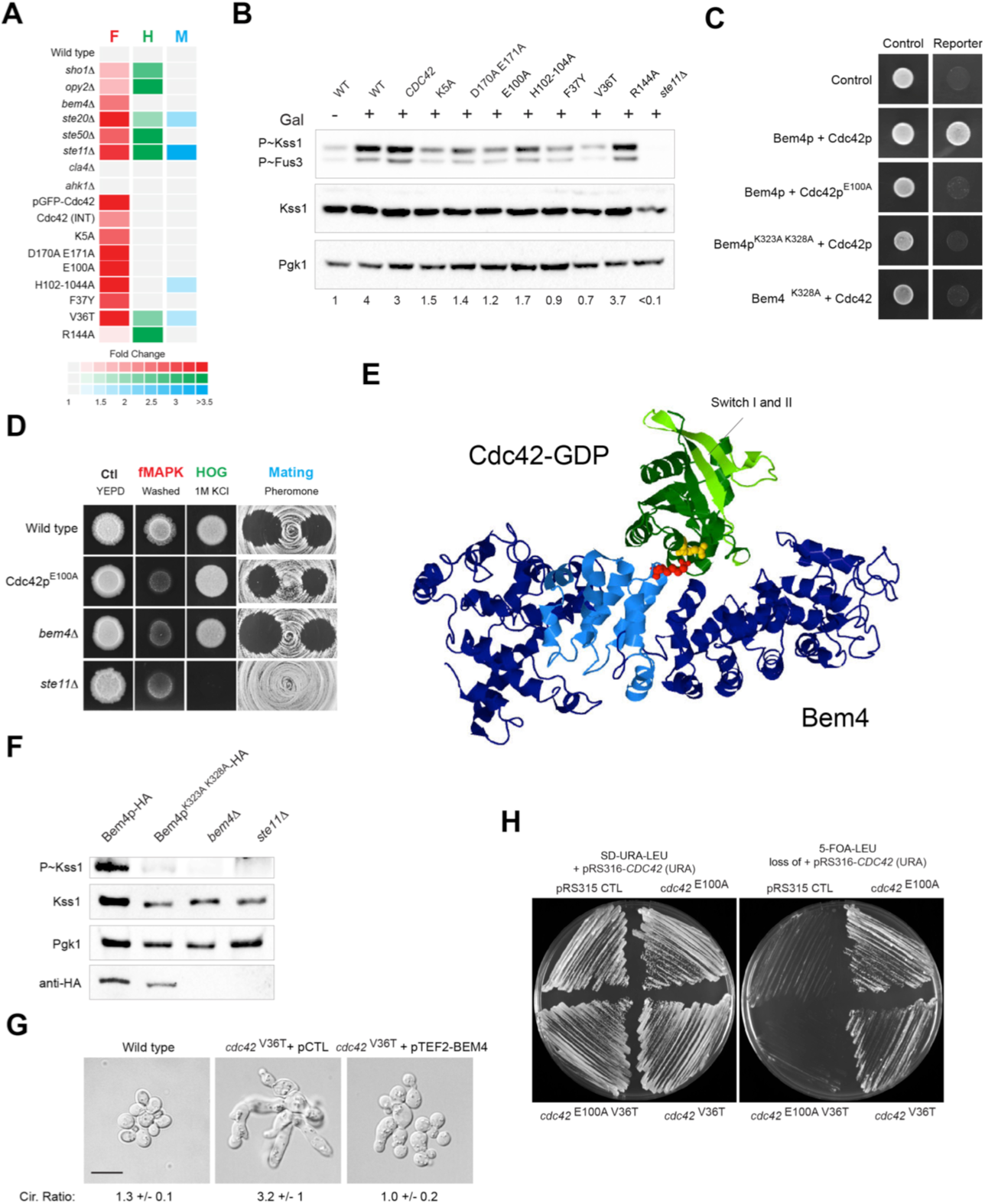
Interaction between Bem4p and Cdc42p is required for fMAPK pathway activity. **A)** *CDC42* alleles that show phenotypes in MAP kinase-dependent assays. Color represents a phenotype in the PWA (fMAPK, F, red), growth on 1M KCl (HOG, H, green) or halo formation in response to α-factor (Mating, M, blue). Color intensity represents the severity of the phenotypic defect based on quantitation by ImageJ analysis; grey, no phenotypic difference from wild type. Controls and several mutants tested are also shown. **B)** Wild-type, a collection of *cdc42* alleles and *ste11*Δ strains were exposed to YEP-GAL for 5.5h to examine P∼Kss1p levels. Numbers indicate the relative band intensity of P∼Kss1p to total Kss1p levels normalized to the control. INT, integrated version of *CDC42*. **C)** Two-hybrid analysis of the indicated versions of Cdc42p and Bem4p. The complete set is available in the supplement (Fig. S8). **D)** Phenotypes of Cdc42p^E100A^ and the *bem4*Δ mutant in tests for MAP kinase pathway activity. **E)** Ribbon diagram of the GDP-bound conformation of Cdc42p, based on the structure of the mammalian Cdc42p and DOCK9 complex (PDB ID#2wmn). The switch I and II regions are highlighted in light green (*54*). The glutamate residue at position 100 in the α3 helix is shown in yellow. The K328 residue in Bem4p is shown in magenta. **F)** P∼Kss1p levels in strains carrying Bem4p^K323A K328A^-HA and controls. Immunoblot shown was also probed with antibodies to Kss1p, Pgk1p and HA. **G)** Microscopic examination of wild-type cells and cells expressing the Cdc42p^V36T^ protein that also contained pTEF2 (pCTL) or pTEF2-*BEM4*. Bar, 10µm. Numbers indicate the circularity index (length-to-width ratio) of cells measured by ImageJ analysis. More than 100 cells were measured for each strain. **H)** The *cdc42::NAT* strain carrying the *pRS316-GFP-CDC42* cover plasmid and indicated *CDC42* alleles integrated at the genomic locus were examined on SD-URA and 5-FOA media to test their requirement for viability.

### Characterization of Mutant Isoforms of Cdc42p with MAPK Pathway-Specific Phenotypes

MAPK pathways were also assessed by tests that were less labor-intensive than phosphoblot analysis and provided a first approximation of MAPK pathway activity. The plate-washing assay (PWA) measures invasive growth and the activity of the fMAPK pathway (*70*). Salt sensitivity was used to measure the activity of the HOG pathway (*71, 72*), and growth arrest by α-factor (halo assays) to measure the activity of the mating pathway (*73*). Data from the functional tests were quantified by ImageJ analysis and color-coded to facilitate interpretation (Fig. 2A). Mutants lacking an intact fMAPK pathway were examined by these tests. As expected, loss of common amplifiers of the fMAPK pathway showed defects in all three pathways (Fig. 2A*, Fig. S3A, ste20*Δ and *ste11*Δ). Also, as expected, several mutants were defective for two of the three pathways (Fig. 2A, Fig. S3A, *sho1*Δ, *opy2*Δ, and *ste50*Δ). Ste50p interacts with Ste11p and is thought to regulate all three pathways (*47, 74–76*) but did not regulate the mating pathway in this context. Loss of the adaptor protein Bem4p only impacted in the fMAPK pathway (Fig. 2A, Fig. S3A, *bem4*Δ). Two proteins that regulate the HOG pathway, the PAK Cla4p (*77, 78*) and the adaptor protein Ahk1p (*79*), were also tested and were not found to regulate the HOG or fMAPK pathways (Fig. 2A, Fig. S3B-F). Thus, Ahk1p and Cla4p do not appear to regulate the fMAPK pathway. These results support and extend our understanding of the proteins that regulate the fMAPK pathway.

Despite the fact that Cdc42p binds to the same PAK, Ste20p, to regulate the three MAPK pathways, versions of Cdc42p have been identified with MAPK pathway-specific phenotypes (*76, 78, 80*). An N-terminal green fluorescent protein (GFP) fusion to Cdc42p, expressed as the sole copy of *CDC42* in the cell, was defective in the fMAPK pathway but not the HOG or mating pathways (Fig. 2A, pGFP-Cdc42, Fig. S4). To follow up on this observation, a collection of *cdc42* alleles that targeted charged, often evolutionarily conserved amino acid residues and functional domains (*81*), was introduced into Σ1278b, which is capable of filamentous growth. Twenty-four non-lethal alleles at 25°C were viable and examined by functional tests (Table S1, Figs. S5-7). Most alleles of *CDC42* did not show a mutant phenotype (Fig. S5-7). However, several alleles of *CDC42* showed pathway-specific phenotypes. Five alleles were mainly defective for the fMAPK pathway based on invasive growth (Fig. 2A, K5A, D170A E171A, E100A, H102-104A, and F37Y) and P∼Kss1p analysis (Fig. 2B). Note that the integrated version of *CDC42* (INT) had a defect in invasive growth. One of the alleles (E100A) has previously been shown to be specifically defective for signaling in the fMAPK pathway (*76, 80*). One allele of *CDC42* was identified that was primarily defective for growth on salt (Fig. 2, A and B, R144A). Several alleles contained mutations in the Switch I domain, which binds PAKs and, as one might expect, were defective for all three pathways (Fig. 2, A and B, V36T).

**Table 1.**
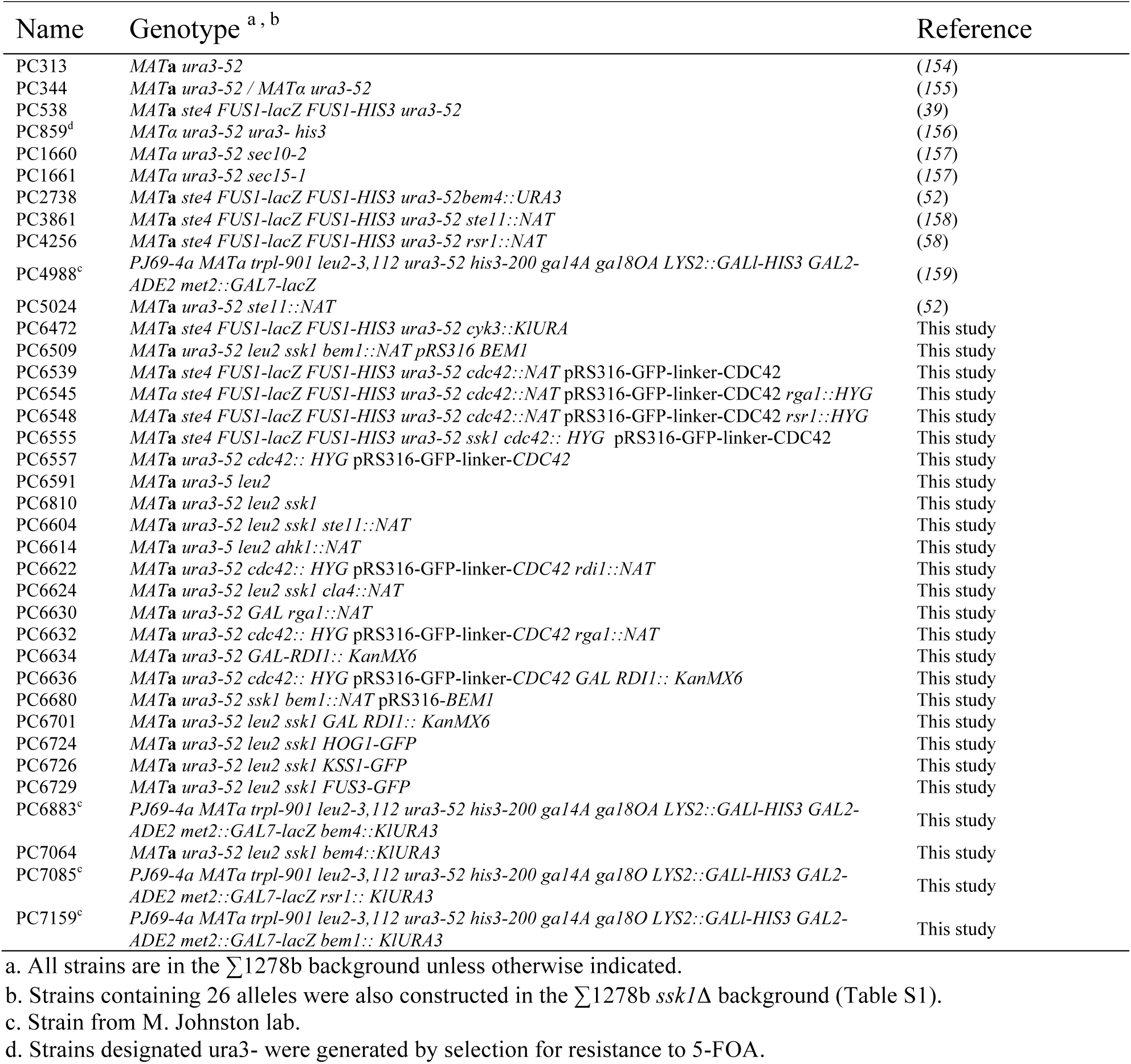
Yeast strains used in the study.

### Interaction Between Bem4p and Cdc42p Stimulates fMAPK Pathway Signaling

Versions of Cdc42p that are specifically defective for fMAPK pathway activity might fail to interact with proteins that regulate the fMAPK pathway. In particular, we considered Bem4p because it is a pathway-specific regulator of the fMAPK pathway [Fig. 2A, (*52*)] and a Cdc42p-interacting protein (*50, 82*). Bem4p interacts with Cdc42p by two-hybrid analysis (*52, 82*). In this study, two-hybrid analysis was examined using a version that mimicked the GDP-bound conformation and did not associate with membranes (*CDC42*^D118A C188S^). By two-hybrid analysis, Bem4p showed reduced interaction with Cdc42p^E100A^ (Fig. 2C). Bem4p was also partially defective for interaction with Cdc42p^K5A^ and Cdc42p^D170A E171A^, but not for interaction with Cdc42p^V36T^, Cdc42p^H102-104A^, or Cdc42p^R144A^ (Fig. S8A). Thus, the signaling defect of Cdc42p^E100A^ in the fMAPK pathway might result from its inability to interact with Bem4p. Consistent with this possibility, the phenotypes of cells carrying Cdc42p^E100A^ phenocopied those of the *bem4*Δ mutant. Like the *bem4*Δ mutant, Cdc42p^E100A^ was defective for invasive growth but not growth on salt or halo formation (Fig. 2D). Cdc42p^E100A^ was also defective for invasive growth by another commonly used test, the single cell invasive growth assay (*33*), but not shmoo formation in response to α-factor (Fig. S7). Similar phenotypes have been reported for the *bem4*Δ mutant (*52*).

Like other monomeric GTPases, Cdc42p undergoes a conformational change upon GTP binding that allows the protein to interact with effector proteins. Most proteins that interact with Cdc42p bind to the active or GTP-bound conformation of the protein. An unusual feature of Bem4p is that it interacts with the GDP-bound (inactive) and GTP-bound (active) conformations of Cdc42p (*52, 82*). We modeled the yeast Cdc42p protein onto the crystal structure of human Cdc42p (*83–85*), which are >80% identical, and found that the glutamic acid at position 100, which is conserved between human and yeast Cdc42p, lies in the third alpha (α3) helix of the protein (Fig. 2E, Cdc42 is in green, E100 is marked in yellow). The α3 helix is on the opposite side of the effector-binding domain of the protein (Fig. 2E, Switch I and II) and is not thought to undergo a conformational change upon nucleotide binding. Therefore, Bem4p may interact with a region of Cdc42p that does not undergo a conformational change. This finding is consistent with its ability to associate with the GDP and GTP-bound conformations of Cdc42p (*52*).

We previously mapped a region of Bem4p from 300 to 400 amino acid (aa) residues as being required for interaction with Cdc42p (*52*). This region exhibits sequence similarity to members of the SmgGDS family of proteins. SmgGDS proteins contain Armadillo (ARM) repeats (*86, 87*) that mediate interactions with Rho GTPases (*86, 88–92*) and other proteins. The predicted structure of Bem4p was determined by bioinformatics algorithms Phyre 2 and iTASSER, which generated similar structures with >90% confidence and theoretical root mean square deviation (iRMSD) values above cutoff. In both structures, Bem4p contained alpha helical motifs that resembled ARM repeats (Fig. 2E, Bem4p is in blue, and the 300-400 region is marked in light blue).

Basic residues in the Bem4p^300-400^ region might mediate electrostatic interactions with the glutamate at position 100 in Cdc42p. Ten basic residues are present in the Bem4p^300-400^ region. Site-directed mutagenesis was used to generate versions of Bem4p that lacked basic residues, based on the predicted minimal free energy between Cdc42p^E100A^ and Bem4p^300-400^ using the protein-docking server Haddock (*93*). Bem4p ^K323A K328A K351A R352A^, Bem4p ^K323A, K328A^, Bem4p ^K351A R352A^, Bem4p ^K323A^, and Bem4p ^K328A^ were constructed and tested. Bem4p ^K323A K328A K351A R352A^, Bem4p ^K323A K328A^, and Bem4p ^K328A^ were defective for interaction with Cdc42p (Fig. 2C, Fig. S8). Bem4p ^K351A R352A^ was partially defective, and Bem4p ^K323A^ was not defective (Fig. S8). The residue K328 in Bem4p had the lowest free energy based on the Haddock algorithm and may be a key residue that mediates interaction with Cdc42p (Fig. 2E, red residue, *Movie 1*). Compensatory mutations did not restore the interaction between Bem4p and Cdc42p (Fig. S8, Bem4p ^K328E^ and Cdc42p^E100A^), which indicates that, like for other Cdc42p-interacting proteins, the interaction between Bem4p and Cdc42p may involve multiple electrostatic contacts. Therefore, basic residues in the ARM-type repeat region of Bem4p are required for interaction with Cdc42p.

Versions of Bem4p that showed reduced interaction with Cdc42p were tested for fMAPK pathway activity. Bem4p-HA ^K323A K328A^ was expressed from the *BEM4* locus at levels similar to the Bem4p-HA protein (Fig. 2F, anti-HA blot). Bem4p-HA ^K323A K328A^ was defective for fMAPK pathway activity based on P∼Kss1p levels (Fig. 2F). Bem4p-HA ^K323A K328A^ was also defective for invasive growth and expression of the *FUS1-HIS3* reporter (Fig. S9), which in strains lacking an intact mating pathway (*ste4*Δ) provides a readout of fMAPK pathway activity (*39, 52, 58, 94, 95*). Bem4p-HA ^K328A^ was also tested but did not show the same defects in MAPK signaling and invasive growth as Bem4p-HA ^K323A K328A^. Thus, both residues of Bem4p are probably important for the regulation of Cdc42p in the fMAPK pathway.

Bem4p can also regulate cell polarity. In particular, high-copy plasmids carrying *BEM4* can suppress the temperature-sensitive growth defects of *cdc42* alleles (*50*). We found that a high-copy plasmid expressing *BEM4* (pTEF2-*BEM4*) suppressed the morphological defect of cells carrying a version of Cdc42p with a single amino acid change in its effector binding domain, Cdc42p^V36T^ (Fig. 2G). To test whether suppression occurred through interaction between Bem4p and Cdc42p, the E100A mutation was introduced into *cdc42* ^V36T^ by site directed mutagenesis. Based on the above results, the E100A mutation would be expected to interfere with the interaction between Bem4p and Cdc42p. We found that cells expressing Cdc42p ^V36T E100A^ had a viability defect compared to cells carrying versions of Cdc42p with either single change, by plasmid-loss experiments using the drug 5-fluoroorotic acid [Fig. 2H, 5-FOA (*96*)). Thus, Cdc42p^V36T^ and Cdc42p^E100A^ may function synergistically to regulate viability. Therefore, the interaction between Bem4p and Cdc42p may also be required for viability when Cdc42p’s interaction with effector proteins, and thus its cell polarity function, is compromised.

### The Polarity Adaptor Bem1p Also Regulates the fMAPK Pathway

Bem4p is a pathway-specific adaptor for the fMAPK pathway. Another adaptor that interacts with Cdc42p, Bem1p, regulates the establishment of polarity (*97–99*). During cell polarization, Bem1p is critical for symmetry breaking by generating positive feedback (*100–103*). Bem1p also regulates the mating (*62, 104*) and HOG pathways (*63*). Bem1p has not been tested for a role in regulating the fMAPK pathway because it is essential for viability in the ∑1278b background but not in laboratory strains, such as S288c (*20*).

To determine whether Bem1p regulates the fMAPK pathway, the *BEM1* gene was deleted in a strain containing the *BEM1* gene on a plasmid (p*BEM1*, provided by D. Lew). Plasmid-loss experiments using 5-FOA showed that the *bem1*Δ mutant was viable under some conditions (YEP-GAL media at 30°C and synthetic media) but not others (YEPD media at 30°C or YEP-GAL media at 25°C). The conditional viability of the *bem1*Δ mutant allowed us to determine whether Bem1p regulates fMAPK. The *bem1*Δ mutant showed reduced fMAPK pathway activity based on phosphorylation of Kss1p (Fig. 3A). The signaling defect of the *bem1*Δ mutant was less severe than the *ste11*Δ mutant and resembled the phenotype of cells lacking other upstream pathway components including Msb2p (*39*), Sho1p (*105*), and Rsr1p (*58*). The *bem1*Δ mutant was also defective for invasive growth by the PWA (Fig. 3B, washed). We also found that the *bem1*Δ mutant had a defect in colony ruffling (Fig. 3B, Magnified), which occurs due to cell-to-cell contacts during biofilm/mat formation, a microbial process that is also controlled by the fMAPK pathway (*106, 107*). The *bem1*Δ mutant was defective in the formation of filamentous chains of cells (Fig. 3B, Scraped, Fig. S10). By the single cell assay, the *bem1*Δ mutant also exhibited polarity and bud-site-selection defects (Fig. 3B, Single cell, Fig. S10). Therefore, Bem1p regulates the fMAPK pathway and is required for invasive growth and biofilm/mat formation.

**Figure 3.**
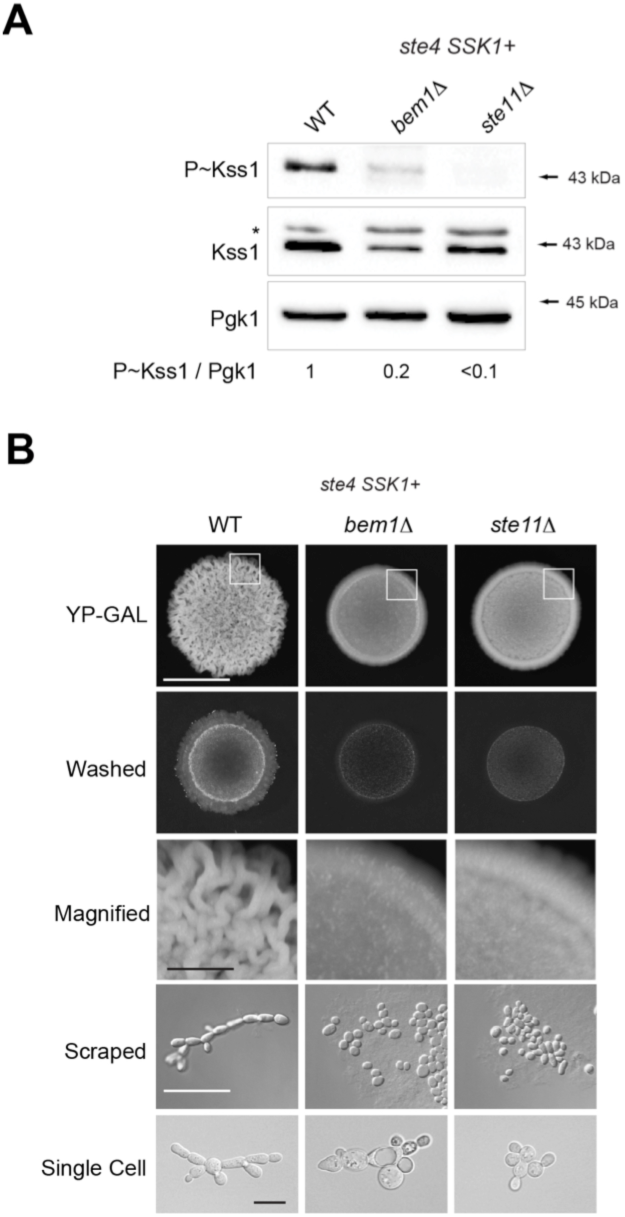
Bem1p regulates the fMAPK pathway. **A)** P∼Kss1p levels in the *bem1*Δ mutant and control cells grown for 5.5h on SD media. Asterisk, background band. Numbers indicate the relative band intensity of P∼Kss1p to total Kss1 levels normalized to the control, which was set to 1. **B)** Analysis of filamentous properties of *bem1*Δ cells by functional tests. Bar, 20 microns.

### Bem1p Regulates the fMAPK Pathway by Recruitment of Ste20p to the Plasma Membrane

Bem1p is composed of domains that perform specific functions during polarity establishment (*99, 108, 109*). These include two SH3 domains, a PX domain, and a PB1 domain (Fig. 4A). Point mutations in these domains selectively cripple individual aspects of Bem1p function (*108*). Versions of Bem1p lacking several domains were tested to ascertain Bem1p’s role in regulating the fMAPK pathway. The fMAPK pathway operates in basal (Fig. 4B, SD-LEU) and activated states (Fig. S11A, YEP-GAL) depending on the condition (*39, 58, 105*). Bem1p^P208L^, which is defective for binding to effector proteins like Ste20p (*62, 101, 110, 111*), was defective for fMAPK pathway signaling (Fig. 4B). Bem1p^P355A^, which prevents an intramolecular interaction that results in a constitutively open conformation of the protein (*108*), was also defective for fMAPK pathway signaling (Fig. 4B). Bem1p^R369A^, which is defective for localization to the plasma membrane and binding to the phosphatidyl inositol polyphosphate (PIP) PI(4, 5)P_2_ at the plasma membrane (*108, 112, 113*), was also defective for fMAPK pathway activity (Fig. 4B). Bem1p^K482A^, which is defective for interaction with Cdc24p (*98, 99, 114-121*), did not show a defect in fMAPK signaling (Fig. 4B) However, a defect was seen in YEP-GAL (Fig. S11A). One interpretation of this result is that Bem1p interacts with effector proteins and Cdc24p at the plasma membrane to regulate the fMAPK pathway. Bem1p may also require cycling between its open and closed conformation to regulate the fMAPK pathway.

**Figure 4.**
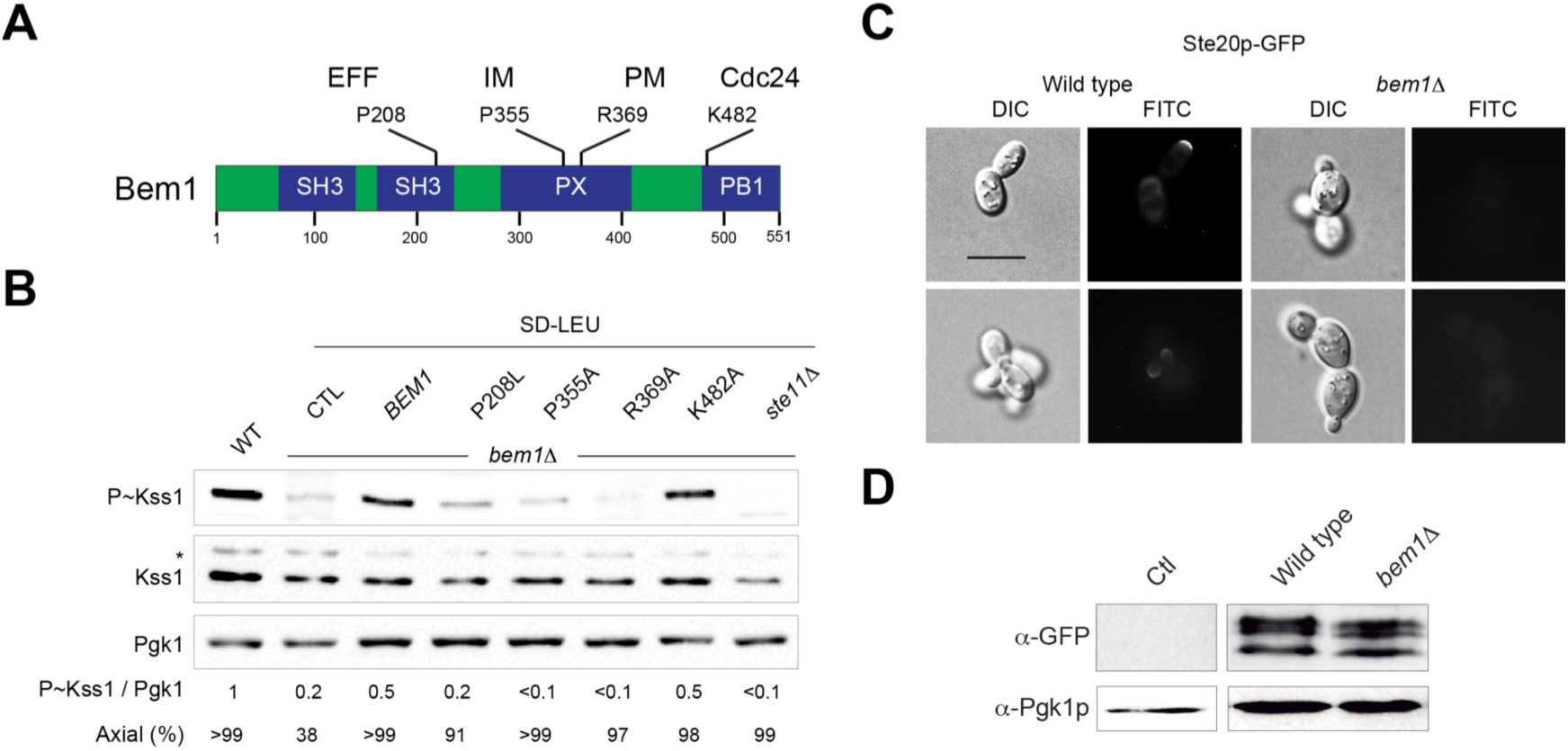
Role of Bem1p in regulating the fMAPK pathway. **A)** Diagram of the Bem1 protein, which shows the effector binding domain (EFF), intramolecular binding domain (IM), plasma membrane (PM) binding domain, and Cdc24p-interacting domain. Alleles used in the study are marked. **B)** Immunoblot analysis of wild-type cells containing wild-type *BEM1* or *BEM1* alleles and control strains grown on SD media. Axial budding expressed as a percentage was determined for cells grown to mid-log phase in SD-LEU medium. Budding pattern was determined by CFW staining. **C)** Localization of Ste20p-GFP in wild-type cells and the *bem1*Δ mutant. Bar, 10 microns. **D)** Immunoblot of Ste20p-GFP in the indicated strains. Antibodies to Pgk1p were used as a control for protein levels.

To test whether Bem1p recruits Ste20p to the plasma membrane, the localization of Ste20p-GFP was examined. In wild-type cells, Ste20p-GFP localized to the cell cortex in small buds and the distal pole in large cells (Fig. 4C, seen in 5-10% of cells). In cells lacking Bem1p, Ste20p-GFP showed a diffuse pattern (Fig. 4C, seen in >200 cells examined). By immunoblot analysis, the level of Ste20p-GFP was the same in wild-type and *bem1*Δ mutant cells (Fig. 4D). This results fits with the fact that Ste20p is the major effector of Cdc42p in the fMAPK pathway (*23, 24, 76*), and with the facts that Bem1p recruits Ste20p to GTP-Cdc42p at the plasma membrane during mating (*23, 24, 76, 110*) and is also required for plasma-membrane localization of Ste20p during filamentous growth (*80*). Therefore, Bem1p promotes plasma membrane localization of Ste20p in the fMAPK pathway.

### Bem1p Regulates Effector Pathways by Different Mechanisms

To compare Bem1p function in Cdc42p-dependent MAPK pathways, phosphoblot and functional assays were performed. As shown above (Figs. 3-4), Bem1p regulated the fMAPK pathway (Fig. 5, A and D). Bem1p did not play a major role in regulating the HOG pathway (Fig. 5B, at least at the 5 min time point shown) although the *bem1*Δ *ssk1*Δ mutant was sensitive to growth on high-osmolarity medium (Fig. 5D, KCl). Bem1p also regulated the mating pathway (Fig. 5, C and D). Therefore, Bem1p is a general regulator of Cdc42p-dependent MAPK pathways in yeast.

**Figure 5.**
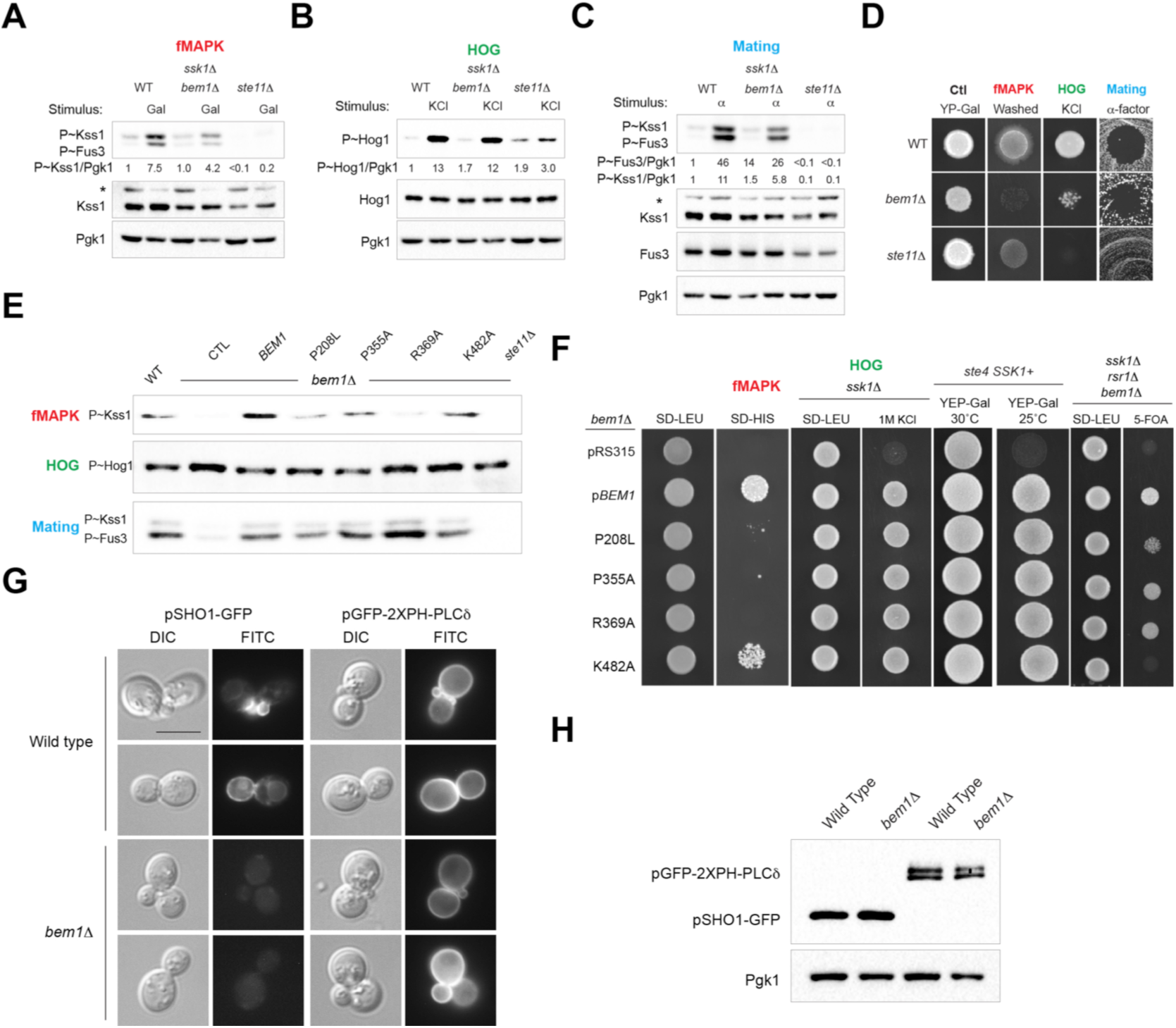
Bem1p regulates different pathways in different ways. **A)** Role of Bem1p in regulating Kss1p phosphorylation in response to galactose. Wild-type and mutant cells were grown for 5.5h in YEP-GAL, specific antibodies were used to detect P∼Kss1p, Kss1p, and Pgk1p. Asterisk denotes a background band. **B)** Role of Bem1p in regulating HOG pathway. P∼Hog1p levels in response to 0.4M KCl (5 min) in the wild-type cells (*ssk1*Δ), *bem1*Δ *ssk1*Δ and *ste11*Δ *ssk1*Δ double mutants. Specific antibodies were alsoused to detect Hog1p and Pgk1p. **C)** Role of Bem1p in response to pheromone. P∼Fus3p and P∼Kss1p levels were examined in response to 6µM of α-factor. Specific antibodies were also used to detect Fus3p, Kss1p and Pgk1p. Asterisk denotes a background band. Numbers indicate the relative band intensity of P∼MAPK to total MAPK levels normalized to the control, which was set to 1. **D)** Wild type, *bem1*Δ and *ste11*Δ (*ssk1*Δ background) cells were spotted on specific media to examine the activity of each MAP kinase pathway. **E)** P∼MAPK levels of *bem1* alleles under specific activating conditions for each MAPK pathway. Cells were grown on YEP-GAL (P∼Kss1p), on YPED + 0.4M KCL (P∼Hog1p) and on YPED + 1.2 of α -factor (P∼Fus3p). **F)** Role of Bem1p in regulating effector pathways. Growth was assessed by spotting *bem1*Δ, *ssk1*Δ *bem1*Δ, or *ssk1*Δ *rsr1*Δ *bem1*Δ cells containing the indicated plasmids onto the indicated media. On SD-HIS, the *FUS1-HIS3* growth reporter was used to examine the activity of the fMAPK pathway. **G)** The localization of Sho1p-GFP and pGFP-2XPH-PLCδ in wild-type cells and the *bem1*Δ mutant. Two examples are shown. **H)** Immunoblot analysis of pGFP-2XPH-PLCδ and Sho1p-GFP in wild-type cells and the *bem1*Δ mutant. Pgk1p immunoblot, control for protein loading.

Does Bem1p regulate Cdc42p-dependent MAPK pathways in the same way? As shown above for basal conditions (Fig. 4), all of the domains of Bem1p tested (P208L, P355A, R369A, and K482A) were required for Bem1p to regulate the fMAPK pathway under activating conditions (Fig. 5E, for the blot plus controls see Fig. S11A). A different pattern was seen in response to salt or pheromone (Fig. 5E, Fig. S11B-C). Therefore, Bem1p regulates Cdc42p-dependent MAPK pathways by different mechanisms. In line with this idea, *bem1* alleles showed different patterns of fMAPK pathway reporter activity (Fig. 5F, SD-HIS) and salt sensitivity (Fig. 5F, KCl). The requirement of Bem1p alleles in production of P∼Fus3p levels did not match that of shmoo formation (Fig. S11D), which might be explained because Bem1p has multiple functions in mating that include direction sensing (*122*) and the formation of mating projections (*119, 120*).

Bem1p is required for viability in the Σ1278b background under some conditions (*20*). The viability defect of the *bem1*Δ mutant was rescued by all versions of Bem1p tested (Fig. 5F, YEP-GAL 25°C). The *bem1*Δ mutant also had a bud-site-selection defect that was rescued by all versions of Bem1p tested (see Fig. 4B, Axial %). Versions of Bem1p that rescued the bud-site-selection defect failed to rescue the fMAPK pathway signaling defect (see Fig. 4B), which addressed the potential concern that Bem1p regulates the fMAPK pathway by its role in regulating bud-site selection (*58*). In some strains, Bem1p requires Rsr1p for viability, which defined a function for Bem1p in symmetry breaking during polarity establishment (*108*). Bem1p also required Rsr1p for viability in the filamentous background (Fig. 5F, 5-FOA). Here, a unique pattern of rescue by *bem1* alleles was seen. The pattern we saw matched the pattern shown in Irazoqui et al. 2003, except that R369A rescued the viability defect in this study but not in Irazoqui et al. 2003. The fact that *bem1* alleles have different phenotypes in symmetry breaking in the two studies might be explained by differences in strain backgrounds. Indeed, Bem1p is required for viability in some strain backgrounds but not others.

Bem1p was also required for the localization of Sho1p-GFP to the plasma membrane (Fig. 5G). This was not due to change in the levels of the protein (Fig. 5H). Sho1p is an integral membrane tetraspan protein that regulates the fMAPK pathway and HOG pathway (*39, 43, 72, 123*). At first, we thought this phenotype might explain how Bem1p regulates the fMAPK pathway. However, all versions of Bem1p tested rescued the localization defect of Sho1p (Fig. S12A). Like Cdc42p (*124, 125*), Bem1p regulates exocytosis, the polarized delivery of vesicles to the plasma membrane (*126, 127*). Bem1p was not required for regulating the distribution of PIPs at the plasma membrane, which might also impact protein localization, based on the localization (Fig. 5G) and levels of a PI(4, 5)P_2_ binding reporter protein [Fig. 5H, pGFP-2XPH-PLCδ (*128*)]. The localization defect of Sho1p-GFP in the *bem1*Δ mutant might result from a defect in exocytosis. Indeed, mutants defective for exocytosis were also defective for the plasma-membrane distribution of Sho1p-GFP (Fig. S12B). Therefore, Bem1p regulates different effector pathways in different ways. In some settings, Bem1p acts as a multifunctional adaptor where one or more domain(s) is required for function. In other settings, Bem1p acts like an inert scaffold, because no single domain of the protein is required.

### Ordering the Activation Sequence of the Cdc42p Module in the fMAPK Pathway

Two adaptors proteins, Bem4p and Bem1p, function at the level of Cdc42p and regulate the fMAPK pathway. As their names suggest (<u>B</u>ud <u>em</u>ergence defect), *BEM1* (*97, 129*) and *BEM4* (*50, 51*) were initially identified as high-copy suppressors of the growth defects of conditional alleles of *cdc24* and/or *cdc42*. Bem1p and Bem4p are not homologs, and the proteins have unrelated amino acid sequences and protein-interaction domains. Loss of either protein results in a defect in the fMAPK pathway (e.g. Fig. 2A, Fig. 3-5). Thus, Bem1p and Bem4p play non-redundant roles in regulating the fMAPK pathway. In support of this idea, a high-copy plasmid containing *BEM1*, which rescues the signaling defect of the *bem1*Δ mutant, did not rescue the fMAPK signaling defect of the *bem4*Δ mutant (Fig. S13). Likewise, a high-copy plasmid containing Bem4p, which rescues the signaling defect of the *bem4*Δ mutant, did not rescue the signaling defect of the *bem1*Δ mutant (Fig. S13). Therefore, Bem4p and Bem1p have non-overlapping functions in regulating the fMAPK pathway.

As adaptor proteins, Bem1p and Bem4p interact with multiple proteins, particularly those that function at the level of the Cdc42p module. Bem1p and Bem4p both interact with Rsr1p, Cdc24p and Cdc42p (*51, 82, 98, 121*). Rsr1p is the Ras-type GTPase that regulates bud-site selection (*99*) and the fMAPK pathway (*58*). Rsr1p can also interact with Cdc42p (*130*). To define how Rsr1p, Bem4p, and Bem1p regulate the fMAPK pathway, genetic suppression analysis was employed. Genetic suppression analysis allows the ordering of components in a pathway by gain- and loss-of-function alleles. GFP-Msb2p (*95*), Sho1p^P120L^ (*131*), and Ste11-4p (*132*) hyper-activate the fMAPK pathway and were tested for bypass of the signaling defects of the *rsr1*Δ, *bem4*Δ, and *bem1*Δ mutants. Hyperactive versions of Msb2p, Sho1p, and Ste11p partially rescued the fMAPK signaling defect of the *rsr1*Δ mutant (Fig. 6, A and D). This finding is consistent with previous results (*58*), which suggest that Msb2p/Sho1p and Rsr1p converge on the Cdc42p module. For the *bem4*Δ mutant, a hyperactive version of Msb2p did not bypass the signaling defect, whereas hyperactive versions of Sho1p and Ste11p did (Fig. 6, B and D). This result indicates that Bem4p functions between Msb2p and Sho1p in the fMAPK pathway. For the *bem1*Δ mutant, hyperactive versions of Msb2p and Sho1p did not bypass the signaling defect, whereas a hyperactive version of Ste11p did (Fig. 6, C and D). This result indicates that Bem1p functions between Sho1p and Ste11p in the fMAPK pathway. Collectively, the data indicate that Bem4p functions upstream of Bem1p in the fMAPK pathway. This conclusion is probably an oversimplification, because the proteins exist in multi-protein complexes and have multiple functions in regulating the fMAPK pathway. However, this order is supported by the fact that Bem4p associates with the GDP-bound conformation of Cdc42p (*52, 82*), whereas Bem1p preferentially associates with the GTP-bound conformation of Cdc42p (*100, 130*). Indeed, two-hybrid analysis showed that the interaction between Bem1p and GDP-Cdc42p required Rsr1p (Fig. 6E, Cdc42 refers to *CDC42*^D118A C188S^), whereas the interaction between Bem4p and GDP-Cdc42p did not require Rsr1p or Bem1p (Fig. 6F).

**Figure 6.**
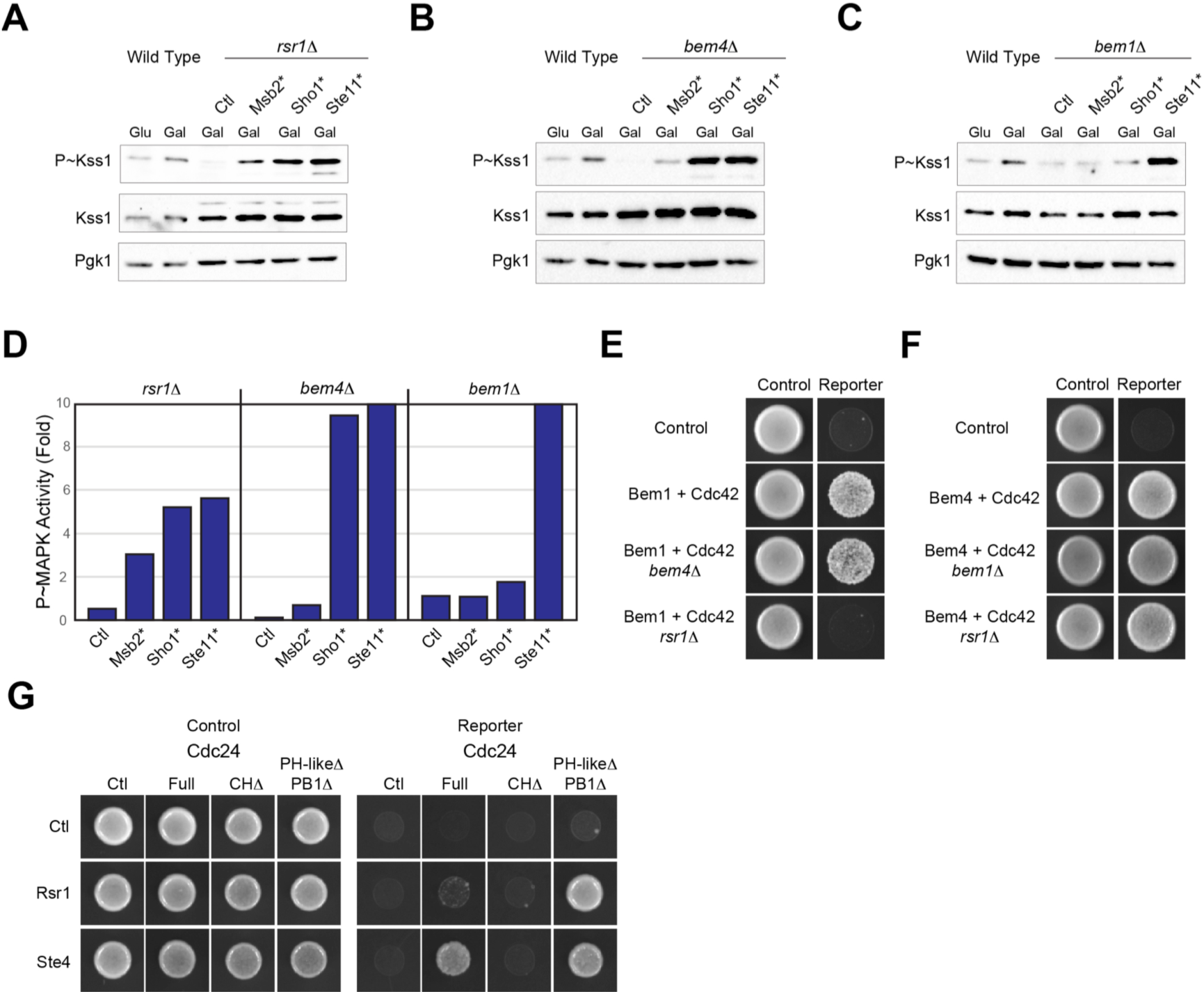
Bem1p and Bem4p regulate the fMAPK pathway at different points in the signaling cascade. **A)** Wild-type cells and the *rsr1*Δ mutant were examined in YEPD (Glu) or YEP-GAL (Gal) media by immunoblot analysis. The *rsr1*Δ mutant contained either a control plasmid (pRS316) or plasmids harboring hyperactive versions of Msb2p (Msb2*, pGFP-MSB2; Sho1*, pSHO1-P120L-GFP, or Ste11*, pSTE11-4). Cells were grown in YEPD or YEP-GAL media for 5.5 h. **B)** Same as in panel A, except the *bem4*Δ mutant was examined. **C)** Same as in panel A, except the *bem1*Δ mutant was examined. **D)** Quantitation of immunoblot data. **E)** Two-hybrid analysis of Bem1p and Cdc42p in wild-type cells and the indicated mutants. **F)** Two-hybrid analysis of Bem4p and Cdc42p in wild-type cells and the indicated mutants. **G)** Two-hybrid analysis of the interactions of Rsr1p and Ste4p with full length Cdc24p and versions that lack the CH (CHΔ) or PH-like and PB1 (PH-likeΔ PB1Δ) domains.

The above results led us to ask how the activation of the fMAPK pathway is initiated at the level of Cdc24p. To address this question, we turned to the mating pathway, which is known to recruit Cdc24p by Ste4p (*133*). By two-hybrid analysis, Ste4p, which regulates the mating pathway and binds the CH domain of Cdc24p, bound to full length Cdc24p, whereas Rsr1p, which regulates the fMAPK pathway and also binds the CH domain of Cdc24p (*134*), did not (Fig. 6G). Rather, Rsr1p bound to a version of Cdc24p lacking an auto-inhibitory domain (PH-likeΔ-PB1Δ). Given that Bem4p binds to the PH-like domain of Cdc24p to relieve auto-inhibition, it is plausible that Ste4p initiates recruitment of Cdc24p in the mating pathway, whereas in the fMAPK pathway, Bem4p has been suggested to bind to Cdc24p to relieve auto-inhibition (*52*), which permits subsequent binding by Rsr1p. Bem1p binds the PB1 domain of Cdc24p and may also initiate recruitment of Cdc24p in the fMAPK pathway. However, Bem1p acts at a later step in the pathway than Bem4p and does not require binding to Cdc24p to activate the fMAPK pathway under some conditions, which argues against this possibility.

## DISCUSSION

Rho GTPases are key regulators of cell polarity and signal transduction in eukaryotes. Many proteins interact with Rho GTPases and impact their functions and activities, including adaptor proteins that link GTPases to their activators and effectors. By examining two adaptors of Cdc42p in yeast, Bem4p and Bem1p, we further define how Cdc42p regulates one of the many pathways in which it functions. Bem1p and Bem4p, as well as Rsr1p, the GTPase that regulates bud-site-selection and the fMAPK pathway, exhibit some degree of pathway selectivity. Bem4p and Rsr1p regulate the fMAPK pathway but not the mating or HOG pathways, while Bem1p regulates the fMAPK pathway by a specific mechanism. Therefore, insights into understanding how a MAP kinase pathway is regulated can help us to understand how Rho GTPases are regulated in a pathway-specific context.

### Cdc42p-Dependent MAPK Pathways Show Different Kinetic Profiles

The way that signaling pathways are activated can shape the output response, as well as signal specificity. MAPK pathway dynamics can be complex, involving fast and slow responses (*135*), oscillations (*136*), and feedback control (*34, 137*). By directly comparing Cdc42p-dependent MAPK pathways in yeast, we show that the Cdc42p-dependent MAPK pathways have different activation profiles. The mating and HOG pathways are rapidly induced to their maximal levels, whereas the fMAPK pathway has slow activation kinetics. The slow activation kinetics of the fMAPK pathway might mean that gene expression is required for MAPK pathway activation. Given that the fMAPK pathway is induced under nutrient-limiting conditions, one possibility is that derepression of glucose-repressed genes (*138*) may be required for fMAPK function. In a related study from our lab (*139*), we show that the fMAPK pathway is regulated throughout the cell cycle and peaks in G_2_/M phase. Cell synchronization experiments also showed that the fMAPK pathway is activated (in G_2_/M) to the same levels as the mating and HOG pathways. Future studies will be required to determine whether the different activation kinetics of Cdc42p-dependent MAPK pathways leads to pathway-specific responses.

### Interaction Between Bem4p and Cdc42p is Critical for fMAPK Pathway Activity

One challenge in studying scaffold-type adaptors that interact with multiple proteins is to define which protein interactions are critical for pathway activity. Bem4p interacts with Cdc24p, Cdc42p, Ste11p, and Kss1p (*52*), yet the critical interactions relevant for Bem4p function have not been explored. Here, we identify a version of Cdc42p (E100A) that is defective for interaction with Bem4p and fMAPK pathway activity. Similarly, specific amino acid residues on Bem4p were identified that are defective for interaction with Cdc42p and fMAPK pathway activity. Together these results define the interaction between Cdc42p and Bem4p as being critical for fMAPK pathway activity. Indeed, this may be the first example in yeast where a protein binding to Cdc42p itself impacts its function in a pathway-specific context.

### A Model for Cdc42p Regulation in the fMAPK Pathway

We also show that the major polarity adaptor of Cdc42p, Bem1p, regulates the fMAPK pathway. Bem1p regulates the fMAPK pathway by recruiting Ste20p, the effector of Cdc42p in the fMAPK pathway, to the plasma membrane. Bem1p regulates the mating pathway through a similar mechanism (*103*). This conclusion satisfies a longstanding problem of how Ste20p is recruited to the plasma membrane during filamentous growth. Recently, Bem1p has been shown to interact with Msb2p in the HOG pathway (*63*). Like Bem1p, Msb2p preferentially associates with the GTP-bound conformation of Cdc42p (*39*). Therefore, Bem1p may function in a protein complex that connects Msb2p, the Cdc42p module, and Ste20p.

Together with previous work, four proteins interact with the Cdc42p module and regulate the fMAPK pathway: Msb2p (*39*), Bem4p (*52*), Rsr1p (*58*), and Bem1p (this study). Each of these proteins has a non-redundant function in the fMAPK pathway. Thus, a current challenge is to define the specific functions for each of these proteins in the regulation of the GTPase module. We have previously shown that Msb2p associates with the GTP-bound conformation of Cdc42p (*39*) and Rsr1p regulates the fMAPK pathway by interacting with the CH domain of Cdc24p (*58*). Here, we show that Bem4p interacts with Cdc42p to regulate the fMAPK pathway, and Bem1p brings Ste20p to the plasma membrane to GTP-Cdc42p. Collectively, these findings allow us to put forward a model for how Cdc42p might be regulated in the fMAPK pathway (Fig. 7).

**Figure 7.**
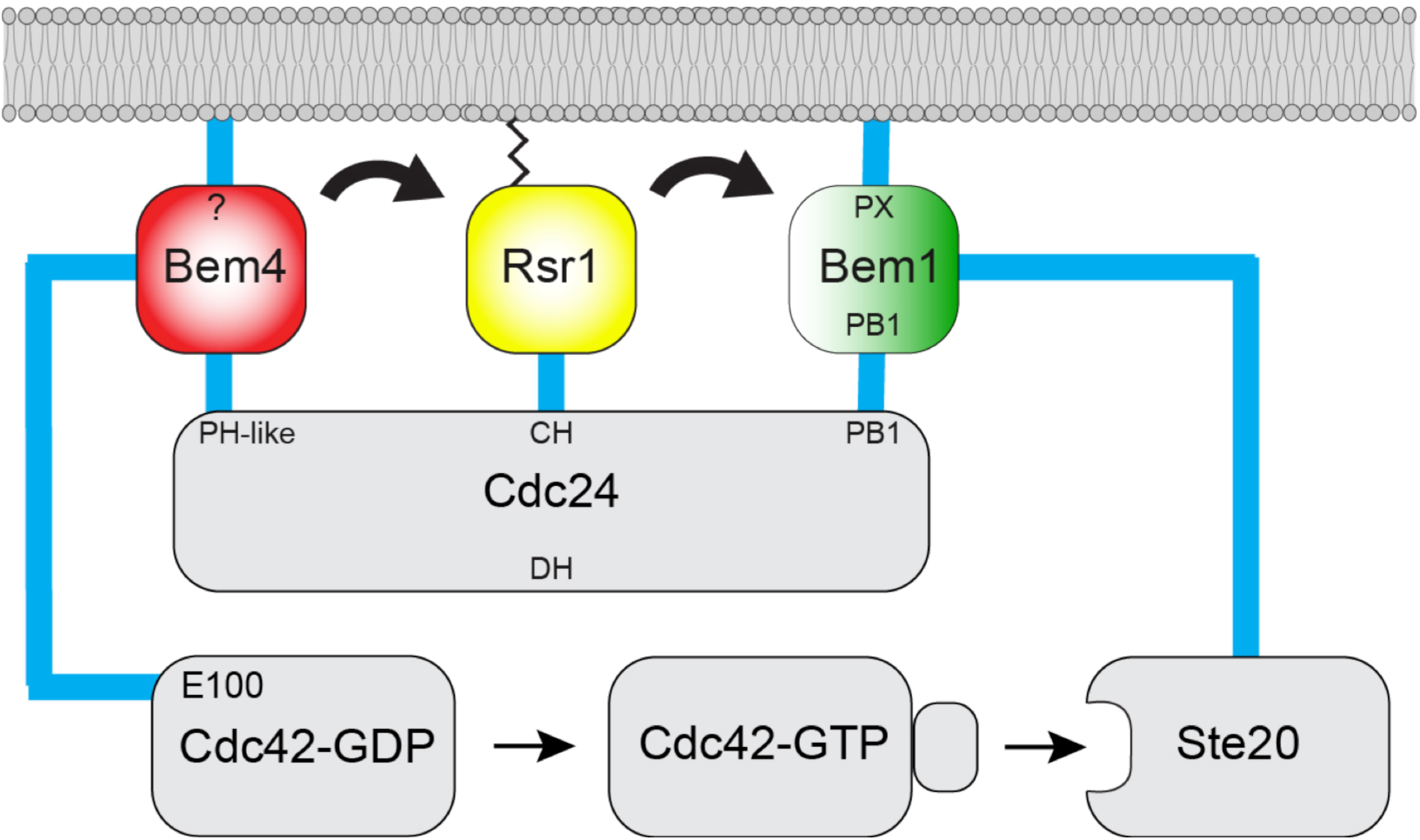
Model for proteins that regulate the Cdc42p module in the fMAPK pathway. Bem4p interacts with the PH-like domain of Cdc24p (*52*) and the α3 helix of Cdc42p (this study) and may initiate recruitment of Cdc24p to the fMAPK pathway. Rsr1p interacts with the CH domain of Cdc24p to regulate the fMAPK pathway (*58*), and Bem1p interacts with the PB1 domain of Cdc24p and with Ste20p to regulate the fMAPK pathway (this study). Each protein is associated with the PM: Rsr1p and Cdc42p by lipid modification (not shown for Cdc42p), Bem1p by the PX domain, and Bem4p in an unknown manner (question mark). Black arrows refer to the order for recruitment and activation of the GTPase module into the fMAPK pathway, which is based on genetic suppression analysis.

One way to begin to define how proteins function in a pathway is by genetic suppression analysis. Genetic suppression analysis using hyperactive versions of fMAPK pathway components showed that Bem4p functions above Bem1p in the fMAPK pathway. This conclusion is consistent with the fact that Bem4p can interact with the inactive (GDP-bound) conformation of Cdc42p, whereas Bem1p primarily interacts with the active (GTP-bound) conformation of the protein. Our results further indicate that Rsr1p does not initiate Cdc24p recruitment to the pathway, because it does not interact with the full-length version of Cdc24p. Bem4p or Bem1p may initiate the recruitment of Cdc24p; however, because the interaction between Bem1p and Cdc24p is not necessary under all conditions for fMAPK pathway activation, Bem4p may initiate the recruitment of Cdc24p to the fMAPK pathway.

It will be interesting to define whether the mechanism surrounding Cdc42p regulation in the fMAPK pathway postulated here is conserved among fungal species. It has previously been shown that Cdc42p and its regulators, Rsr1p and Bem1p, contribute to hyphal formation and virulence in the major human fungal pathogen *Candida albicans* (*140*). Perhaps Rsr1p, Bem1p, and Bem4p regulate the orthologous MAP kinase pathway in *C. albicans* and other pathogens.

### Comparison of Cdc42p Regulation in the Mating and fMAPK Pathways

Proteins that regulate Cdc42p in the fMAPK pathway characterized in this study can be compared to proteins that regulate Cdc42p in the mating pathway, which has been characterized previously (*73, 141, 142*). A potentially analogous activation mechanism comes from comparing Ste4p, the ß-subunit of the heterotrimeric G-protein in the mating pathway, to Rsr1p in the fMAPK pathway. Both proteins interact with the CH domain of Cdc24p (*58, 133*), and both proteins are tied to the plasma membrane. Ste4p is anchored to the plasma membrane by Ste18p, which is a palmitoylated and farnesylated protein (*100, 133, 143-146*), and Rsr1p is modified by a lipid moiety (*147*). Thus, Cdc24p may be activated in the same way by Ste4p in the mating pathway, and Rsr1p in the fMAPK pathway. Ste4p can bind to full-length Cdc24p, perhaps to initiate signaling, whereas Rsr1p binds to versions of Cdc24p that lack auto-inhibitory domains. During mating, Ste4p may be critical for initiating recruitment of Cdc24p into the mating pathway, whereas during filamentous growth Rsr1p is not likely to initiate Cdc24p recruitment to the fMAPK pathway.

During mating, the interaction between Ste4p and Cdc24p is stabilized by Far1p, an adaptor protein in the mating pathway that is also required for cell-cycle arrest in response to pheromone (*133, 143, 148*). Bem4p may play an analogous role in the fMAPK pathway. Bem4p interacts with Rsr1p (*82*) and Cdc24p (*68*) and might stabilize the Rsr1p-Cdc24p interaction. Bem4p also interacts with Cdc42p to regulate the fMAPK pathway. Finally, Bem1p regulates the mating and fMAPK pathways by plasma-membrane recruitment of Ste20p (Fig. 7), although this may be an oversimplification, as there are differences in the ways that Bem1p regulates the two pathways. Therefore, the mating and fMAPK pathways utilize different proteins to regulate the same GTPase module.

### Bem1p Regulates Different Pathways in Different Ways

We also found that Bem1p regulates different effector pathways in different ways. Bem1p regulates multiple Cdc42p-dependent processes, including polarity establishment, exocytosis, and MAPK signaling. Remarkably, different domains of Bem1p play different roles in regulating these different pathways. In some pathways, Bem1p acted as a multifunctional adaptor because multiple domains were required for Bem1p function. In other pathways, Bem1p functioned as an inert scaffold, as none of the functional domains of the Bem1p protein tested were required. Two general conclusions can be reached from these observations. First, context is critically important for studying adaptor function. Assessing Bem1p function was sensitive to the particular strain, assay, and pathway being tested. The second conclusion is that adaptors may operate in different ways in different contexts.

## ABBREVIATIONS

ATA, 3-AT, 3-Amino-1,2,4 triazole; 5-FOA, 5-fluoroorotic acid; aa, amino acid; CFW, Calcofluor White; DIC, differential-interference contrast; GAP, GTPase activating protein; GEF, guanine nucleotide exchange factor; HOG, high osmolarity glycerol response; MAPK, mitogen activated protein kinase; PAK, p21 activated kinase; PM, plasma membrane; and PIP, phosphatidylinositol phosphate; and Rho, Ras homology.

## ACKNOWLEDGEMENTS

Thanks to Daniel Lew (Duke University), Rong Li (Johns Hopkins University), John Pringle (Stanford University, Palo Alto, CA), Haruo Saito (University of Tokyo, Japan), David Drubin (UC Berkeley), Peter Novick (University of California, San Diego, CA), Stan Fields (University of Washington), Scott Emr (Cornell University), and Hiten Madhani (University of California San Francisco) for reagents. Thanks to Lawrence Kelley (Imperial College, London) for help with modeling predictions. Nadia Vadaie, Lauren Caccamise, and Colin Chavel helped with experiments. Atindra Pujari provided editorial assistance with the manuscript. The work was supported by a grant from the NIH (GM098629).

## MATERIALS AND METHODS

### Media and Growth Conditions

Experiments were performed at 30°C unless otherwise indicated. Yeast extract, peptone, and dextrose (YEPD) [2% glucose (D)], YEP-GAL (2% GAL) and synthetic complete media containing yeast nitrogen and 2% Glucose (SD) or 2% galactose (S-GAL) were used. Amino acids were added to the synthetic media as required. For some experiments, 0.2% glucose was used as indicated.

### Strains and Plasmids

Strains are listed in Table 1 and plasmids in Table 2. *Escherichia coli* and *Saccharomyces cerevisiae* strains were manipulated using standard methods (*149*). Gene disruptions and *GAL1* promoter fusions were made by PCR-based methods (*150*). Some gene deletions were constructed using cassettes that contained antibiotic resistance markers (*151*). Gene disruptions were confirmed by PCR Southern analysis and phenotype.

**Table 2.** Plasmids used in the study.

| Name | Description | Reference |
| --- | --- | --- |
| PC1405 | pFRE-lacZ | (28) |
| PC1422 | pRS315 | (152) |
| PC1441 | YCp50-STE11-4 | (Stevenson et al. 1992) |
| PC1601 | pRS316-SHO1-GFP | (160) |
| PC1614 | pRS316-SHO1(D16H)-GFP | (160) |
| PC1696 | pGFP-MSB2 | (161) |
| PC1715 | pRS316-SHO1(P120L)-GFP | (161) |
| PC1882 | pRS316-GAL-SHO1(D16H)-GFP::KanMX6 | (161) |
| PC2205 | pNAT | (162) |
| PC2206 | pHYG | (162) |
| PC2207 | pRS316 | (152) |
| PC2560 | pRS426 GFP-2XPH-PLCδ | (128) |
| PC4190 | pOAD-BEM4 | (163) |
| PC4205 | pOAD | (163) |
| PC4206 | pOBD-2 | (163) |
| PC4232 | pOBD-2 CDC42(D57Y, C188S) | (52) |
| PC4590 | pOAD-RSR1 | This study |
| PC4878 | pV84-FUS1-lacZ | This study |
| PC6025 | pYM111 pRS416 8XCRE-LacZ Amp Ura | (45) |
| PC6367 | YE p352-TEF2-BEM4-GFP | (52) |
| PC6455 | pRS306-GFP-linker-CDC42 (pDLB3609) | (108) |
| PC6454 | pRS316-GFP-linker-CDC42 | This Study |
| PC6514 | BEM1p-BEM1-12XMYC-SWE1t (pDLB2374) | (108) |
| PC6515 | BEM1 p-bem1P208L-12XMYC-SWE1t (pDLB2375) | (108) |
| PC6516 | BEM1 p-bem1P355A-12XMYC-SWE1t (pDLB2377) | (108) |
| PC6517 | BEM1p-bem1R369A-12XMYC-SWE1t (pDLB2378) | (108) |
| PC6518 | BEM1p-bem1K482A-12XMYC-SWE1t (pDLB2379) | (108) |
| PC6892 | 2μ SEC4: URA3; pNB142 | (164) |
| PC6893 | 2μ SEC15: URA3; pNB192 | (164) |
| PC6894 | EXO70, 2u, URA3 | (164) |
| PC6519 | pRS316-BEM1 | D. Lew |
| PC6797 | pOBD-2-CDC42(K5A, D57Y, C188S) | This study |
| PC6798 | pOBD-2-CDC42(R144A, D57Y, C188S) | This study |
| PC6799 | pOBD-2-CDC42(E100A, D57Y, C188S) | This study |
| PC6800 | pOBD-2-CDC42(V36T, D57Y, C188S) | This study |
| PC6802 | pOBD-2-CDC42(H102A,H103A, H104A D57Y, C188S) | This study |
| PC6805 | pOBD2-CDC42(F37Y, D57Y, C188S) | This study |
| PC6831 | pOAD-BEM1 | This study |
| PC6833 | pOBD-2 CDC42(C188S) | This study |
| PC6834 | pOBD-2 CDC42(C188S, G12V) | This study |
| PC6835 | pOBD-2 CDC42(C188S, T17N) | This study |
| PC6836 | pOBD-2 CDC42(C188S, D118A) | This study |
| PC7025 | pRS316-HIS-CDC42 | This study |
| PC7028 | pYIp352-TEF2-RDI1 | This study |
| PC7096 | pOAD-BEM4(K323A, K328A, K351A, R352A) | This study |
| PC7097 | pOAD-BEM4(K351A, R352A) | This study |
| PC7112 | pOAD-BEM4(K323A,K328A) | This study |
| PC7160 | pOAD-BEM4(K323A) | This study |
| PC7161 | pOAD-BEM4(K328A) | This study |
| PC7162 | pOAD-BEM4(K328E) | This study |
| PC7163 | pOBD-CDC42(E100K, D57Y, C188S) | This study |

Plasmids pRS315 and pRS316 have been described (*152*). Plasmids carrying *BEM1p-BEM1-12XMYC-SWE1t* (pDLB2374), *BEM1p-bem1*^P208L^*-12XMYC-SWE1t* (pDLB2375), *BEM1p-bem1*^P355A^*-12XMYC-SWE1t* (pDLB2377), *BEM1p-bem1*^R369A^*-12XMYC-SWE1t* (pDLB2378), *BEM1p-bem1*^K482A^*-12XMYC-SWE1t* (pDLB2379) have been described (*108*). These plasmids and pRS316-*BEM1* were generously provided by Dr. Daniel Lew at Duke University School of Medicine.

To generate PC6810, the *LEU2* gene was disrupted by *NAT* in PC313 by homologous recombination. The NAT gene was replaced with *URA3* cassette (HA-URA3-HA). The strain was treated on 5-FOA to force out the *URA3* selection marker. In this strain, the *SSK1* gene was disrupted using the *NAT* gene, which was again replaced with *URA3* cassette. The resulting strain PC6810 is a derivative of PC313 which is *MAT***a** *ura3-52* carrying unmarked *ssk1* and *leu2* gene deletions.

To generate pRS316-GFP-Linker-Cdc42p, the insert was subcloned from pDLB3609 (*108*) using EcoR1 (NEB Cat # R0101, Ipswich, MA) and Sal1 (NEB Cat # R3138, Ipswich, MA). To generate pRS315-GFP-Linker-Cdc42p the insert was subcloned from pDLB 3609 (*108*) using PstI (NEB Cat # R3140, Ipswich, MA) and SalI (NEB Cat # R3138, Ipswich, MA). To generate pYEp352-*TEF2*-*RDI1* (PC7028), the *RDI1* gene was amplified from the genomic DNA cloned in plasmid carrying *TEF2* promoter using Xba1 (NEB Cat # R0145, Ipswich, MA) and Sal1 (NEB Cat # R3138, Ipswich, MA). To create the Bem4p alleles for two-hybrid analysis PC4190 with pOAD *BEM4* was utilized. pV84-*FUS1-lacZ* was constructed in plasmid V84 by homologous recombination.

### Functional Tests to Evaluate Budding Pattern and MAP Kinase Activity

The budding pattern was determined as described (*58*). Budding pattern was determined by bud position and bud scar position to the proximal, equatorial, or distal parts of the cell. For some experiments, cells were grown to mid-log phase and resuspended in 1ml of water. Cells were stained with Fluorescent brightener 28 (Calcofluor White, CFW) (Sigma-Aldrich Life Science and Biochemicals, St. Louis, MO) using a final concentration of 0.01% for 10 min, at 25°C. Cells were washed one time in water and examined by microscopy.

The PWA was performed as described (*70*). The single cell invasive growth assay was performed as described (*33*). To evaluate the HOG pathway, cells were spotted onto media supplemented with 0.9 M KCl. Halo assays were performed as described (*73*). Cells were grown for 16 h to saturation. Cell density was measured by adding 20 µl overnight culture to 180 µl water (OD_600nm_ ∼ 0.2). Approximately 5 µl overnight culture was added to 300 µl water and top spread on agar plates. 3µl and 10µl alpha factor (1mg/ml) was added on the dried plates and incubated at 30°C for 2 d. ImageJ analysis was used to quantitate colony growth or the degree of agar invasion by measuring signal intensity compared to background.

Experiments to evaluate the morphogenetic response to pheromone (shmoos) were performed by microscopy. The *FUS1-HIS3* reporter in cells lacking *STE4* was used to evaluate the activity of the fMAPK pathway (*39, 52, 58, 94, 95*) and was measured by spotting cells onto SD-HIS or SD-HIS medium with 3-Amino-1,2,4 triazole (ATA). Tests for viability were performed by plasmid loss experiments of the *URA3* gene on 5-fluoroorotic acid [5-FOA, (*96*)].

### Construction and Evaluation of a Collection of cdc42 Alleles in the ∑1278b Background

A collection of *CDC42* alleles that was previously generated (*81*) was introduced into a filamentous (∑1278b) strain lacking *SSK1* (PC6810). Plasmids from *E. coli* strains harboring *CDC42* alleles linked to the *LEU2* selection marker and flanking sequence required for homologous recombination were digested with BanII (NEB Cat # R0119S, Ipswich, MA) and Xba1 (NEB# R0145S, Ipswich, MA) and transformed into yeast lacking a genomic copy of *CDC42* (cdc42::*NAT*) and containing *pRS316 GFP-CDC42* (PC6454). Selection for *LEU2+/NAT –* colonies favored integration of the alleles at the *CDC42* locus. Selection on 5-FOA was used to force out the p*GFP*-*CDC42* plasmid and determine whether the alleles were viable. To confirm the DNA sequence of each allele, genomic DNA was extracted. The *CDC42* gene and flanking region was amplified by polymerase chain reaction (PCR). PCR products were purified and the entire *CDC42* gene was evaluated by DNA sequencing analysis (Roswell Park Cancer Research Center, Buffalo, NY). In addition to the mutations mentioned in Table S1, the library also carries a polymorphic variation at A190T. Twenty-four of forty alleles were obtained by this method that were viable without the cover plasmid (*pRS316-GFP-CDC42*). Seven alleles were unable to lose the p*GFP*-*CDC42* plasmid, indicating that they produced versions of Cdc42p that compromised its essential function, and one allele was not obtained. Introduction of the wild-type version of *CDC42* resulted in an invasive growth defect that was taken into account when examining the other alleles.

### β-Galactosidase Assays

Wild-type cells (PC313), or cells lacking the redundant branch of the HOG pathway (*ssk1*Δ PC6810) were transformed with plasmids *pFRE-lacZ/URA3* 2µ AMP (PC1405) provided by H. Madhani (*28*), pRS416 *8XCRE-LacZ* AMP URA3 (pYM111 or PC6025) provided by Haruo Saito (*78*), or pV84-*FUS1-lacZ* (PC4878, this study). Cells harboring plasmids were grown to saturation at 30°C in SD-URA medium to maintain selection for the plasmids. Cells in mid-log phase in YEPD were collected, washed three times in distilled water, and added to YEP-GAL media, YEPD medium supplemented with 0.4M KCl, or YEPD medium supplemented with 1.2 µM alpha-factor [5 µl of 1mg/ml α-factor (0.6 µM) in 5 ml, unless specified]. One milliliter of culture aliquots was collected at the indicated time points, assessed for cell density at A_600_, and stored at −80°C.

β-galactosidase assays were performed as described (*39*). Cells were resuspended in 100 µl Z-buffer [44.32 ml H_2_O with 5 ml phosphate buffer (0.6 M Na_2_HPO_4_ + 0.4 M NaH_2_PO_4_), 0.5 ml 1M KCl, 50 µl 1M MgSO_4_, 135 µl ß-mercaptoethanol (BME)] containing 2 µl of 5% sarkosyl and 2 µl toluene. Tubes were incubated at 37°C for 30 min. 650 µl of Z-buffer containing ortho-Nitrophenyl-β-galactoside (ONPG) was added to each tube. Reactions were stopped with 250 µl of 1M Na_2_CO_3_, and the time was recorded. Cell extracts were removed by centrifugation (13,000 rpm for 3 mins), and 200 µl of the supernatant was used to determine optical density at A_420_. Miller Units were calculated as 1000 X A_420_/(A_600_ X time). Experiments were performed in three independent replicates. The two-tailed paired student’s t-test was used to measure the statistical significance between samples from multiple trials using ProStat (Poly Software International, Inc.).

### Phosphoblot and Immunoblot Analysis

Cells were induced for immunoblots according to standard conditions and protocols. For basal conditions, 750 µl of cells grown for 16 h were inoculated into fresh YEPD media (15 ml) and incubated for 3 h to mid-log phase. For inductions, mid-log phase cultures were harvested, washed three times with water, and resuspended in YEP-Gal (15 ml), YEPD + 0.4 M KCl (15 ml), or YEPD + α-factor (15 ml) media over a designated time series.

Immunoblots were performed as described (*58*) with the following exceptions in antibody dilution and blocking buffer. The p44/p42 antibodies (Cell Signaling Technology, Danvers, MA; Cat #4370) and Phospho p38 antibodies (Cell Signaling Technologies Danvers MA #9211) were used to detect P∼MAP kinases and diluted 1:10,000 in 5% BSA. Anti-Kss1p antibodies (Santa Cruz Biotechnology, Santa Cruz, CA; Cat #6775) and anti-Hog1p antibodies (Santa Cruz Biotechnology, Santa Cruz CA; #yC-20) were used at dilutions 1:10000 and 1:5000, respectively, in 5% nonfat dried milk. Anti-Fus3p antiserum (Santa Cruz Biotechnology, Santa Cruz, CA; #6773) was used at a 1:10000 dilution in 5% BSA. Anti-Pgk1p antibodies (Life Technologies; Camarillo, CA; Cat #459250) were used at 1:10,000 dilution. Secondary antibodies goat anti-mouse IgG–HRP (Bio-Rad Laboratories, Hercules, CA; Cat #170-6516), goat anti-rabbit IgG-HRP (Jackson ImmunoResearch Laboratories, Inc., West Grove, PA; Cat #111-035-144), and donkey anti-goat IgG-HRP (Santa Cruz Biotechnology, Santa Cruz, CA; Cat #sc-2020) were used. Quantitation of band intensities for immunoblot analysis was performed with Image Lab Software (Bio-Rad, Inc.) at an exposure with no saturated pixel in any lane. Background subtraction was performed according to the software user guide provided by the manufacturer. Band intensities of phosphoproteins were measured against total protein levels based on Pgk1p band intensity.

### Site-Directed Mutagenesis

To insert point mutations in plasmids carrying the *CDC42* or *BEM4* genes, the GeneArt™ Site-Directed Mutagenesis Kit was used (Thermo Fisher Cat#A13282, Grand Island, New York) according to the manufacturer’s protocol. Briefly, an overlapping ∼40 nucleotide primer set was designed with the desired point mutations designed in the center of the primers. The template was amplified using AccuPrime™ Pfx DNA polymerase (Thermo Fisher Cat#12344024, Grand Island, New York). The template was methylated by adding DNA methylase and S-Adenosyl methionine in the PCR reaction, which was provided by the manufacturer. Following PCR, the free ends of the PCR product were recombined *in vitro* by the addition of recombinase enzyme mix. The recombined PCR product was transformed into *E. coli* cells (One Shot® MAX Efficiency® DH5α™-T1^R^). The specially designed host *E. coli* strain that contains *Mcr*Bc endonuclease digested the methylated template, leaving the non-methylated PCR amplified mutated product intact.

### DIC and Fluorescence Microscopy

Differential-interference contrast (DIC) and fluorescence microscopy of cells were performed using DIC and fluorescence filter sets on an Axioplan 2 fluorescent microscope (Zeiss) with a PLAN-APOCHROMAT 100X/1.4 (oil) objective (N.A. 0.17)(Zeiss). For most experiments, proteins were visualized by resuspending cells in water at 25°C. Digital images were obtained with the Axiocam MRm camera (Zeiss). Axiovision 4.4 software (Zeiss) was used for image acquisition and analysis.

### Protein Modeling

To visualize the putative structure of Bem4p, two web based protein fold recognition and function prediction servers were utilized: Phyre2 (http://www.sbg.bio.ic.ac.uk/phyre2/html/page.cgi?id=index) and iTASSER (https://zhanglab.ccmb.med.umich.edu/I-TASSER/). Phyre2 is designed to predict structure based on primary amino acid sequence (*153*) and was used to model the structure of yeast Cdc42p and alleles using information from the crystal structure of *Homo sapiens Cdc42p* in Protein Data Bank (PDB https://www.rcsb.org/pdb/home/home.do). The protein-docking server Haddock (*93*) was also used and is publically available (http://milou.science.uu.nl/services/HADDOCK2.2/).

## SUPPLEMENTAL MATERIAL

### SUPPLEMENTAL FIGURE LEGENDS

**Figure S1.**
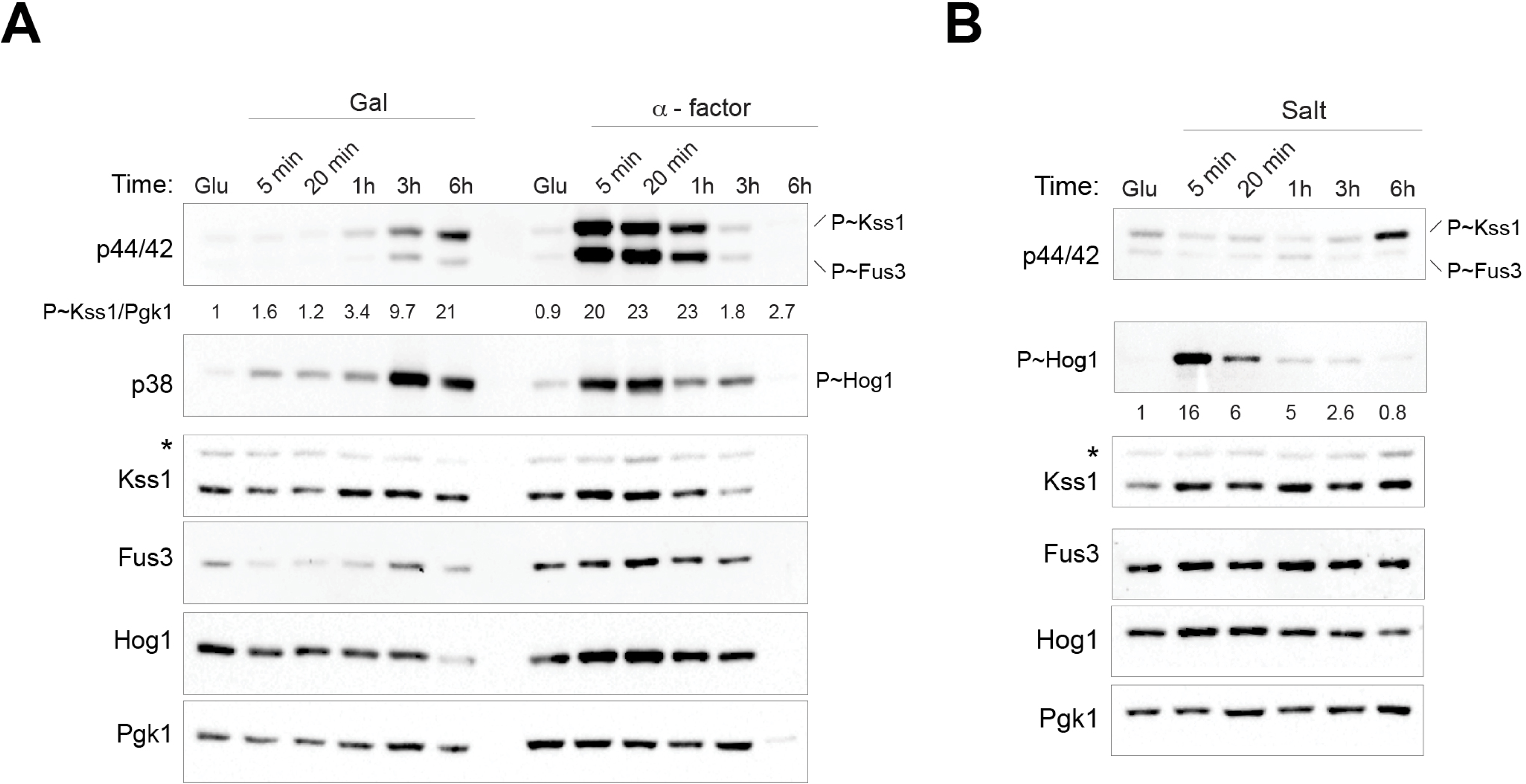
Analysis of the phosphorylated (P∼) MAP kinases. **A)** Immunoblots were probed with p42/p44 antibodies to detect P∼Fus3p and P∼Kss1p. Wild-type cells were grown in YEP-GAL (Gal) or YEPD (Glu) with α-factor during the indicated time points. Immunoblots were also probed with antibodies specific for Fus3p, Kss1p, and Hog1p. Anti-Pgk1p antibodies were used as a control for protein levels. Numbers indicate relative band intensity of P∼MAPK to total MAPK levels normalized to the control condition, which was set to 1. **B)** Wild-type cells were grown in YEPD (Glu) or YEPD supplemented with 0.4 M KCl, p38 antibodies were used to detect P∼Hog1p, the results were analyzed as in panel A. Asterisk, background band.

**Figure S2.**
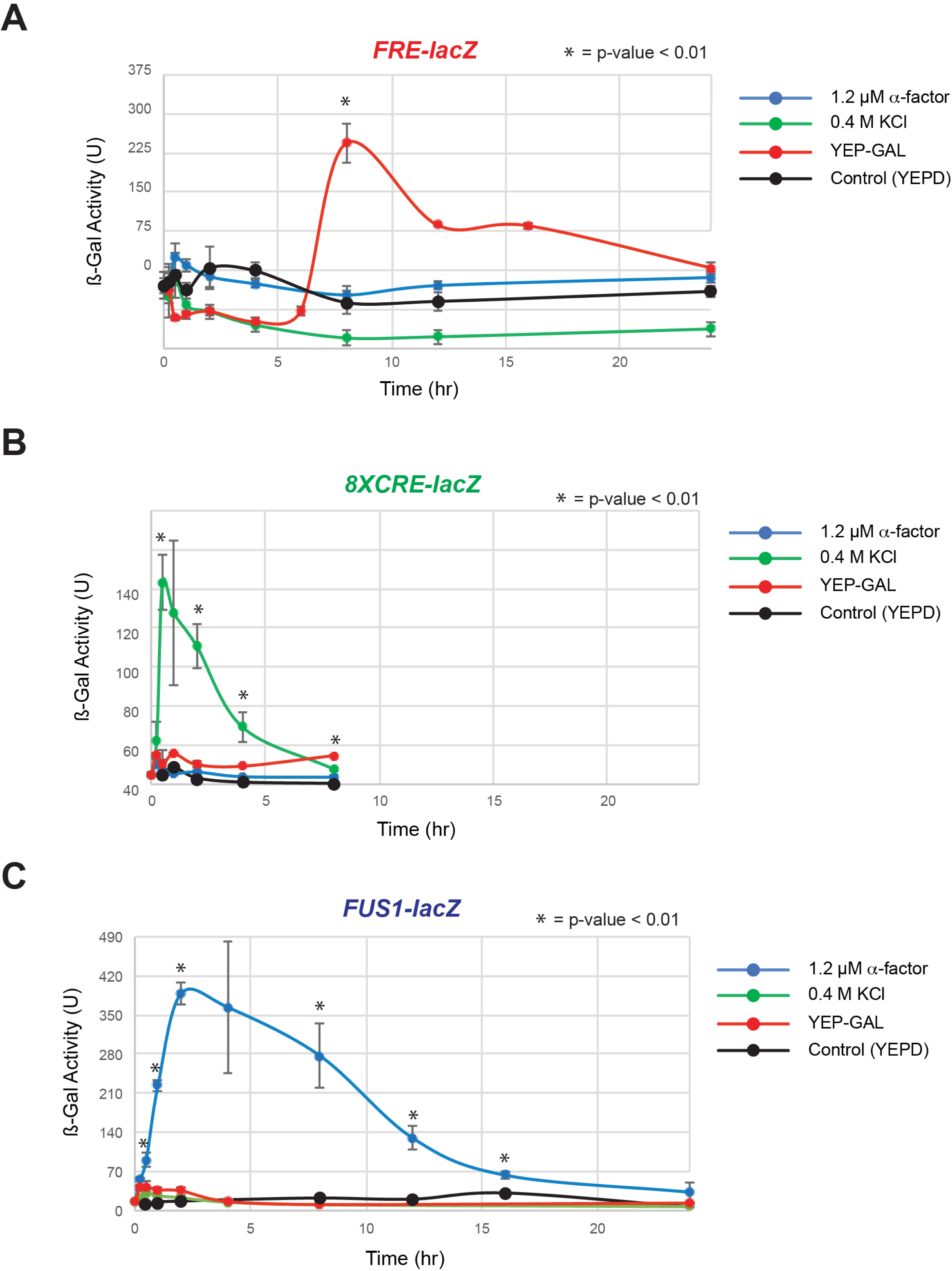
β-galactosidase activities of transcriptional reporters for the indicated MAPK pathways that share components. **A)** Cells (PC313) carrying the mating pathway *FUS1-lacZ* reporter were grown to mid-log phase in YEPD (grey) or induced with 6 µM α-factor (blue), 0.4 M KCl (green), or transferred to YEP-Gal (red) for the indicated times. Data points for the α-factor inductions at 0.5 h, 1 h, and 2 h were omitted to emphasize the levels of activity of the other pathways. Asterisk, p-value < 0.01, with respect to YEPD values. N.S., not significant. **B)** The activity of the HOG reporter, *p8XCRE-lacZ*, was examined as described in panel S2A. Data points for the salt inductions at 0.5 h, 1 h, and 2 h were omitted to emphasize the activity of the other pathways. **C)** The activity of the fMAPK reporter, *pFRE-lacZ*, was examined as described in panel S2A.

**Figure S3.**
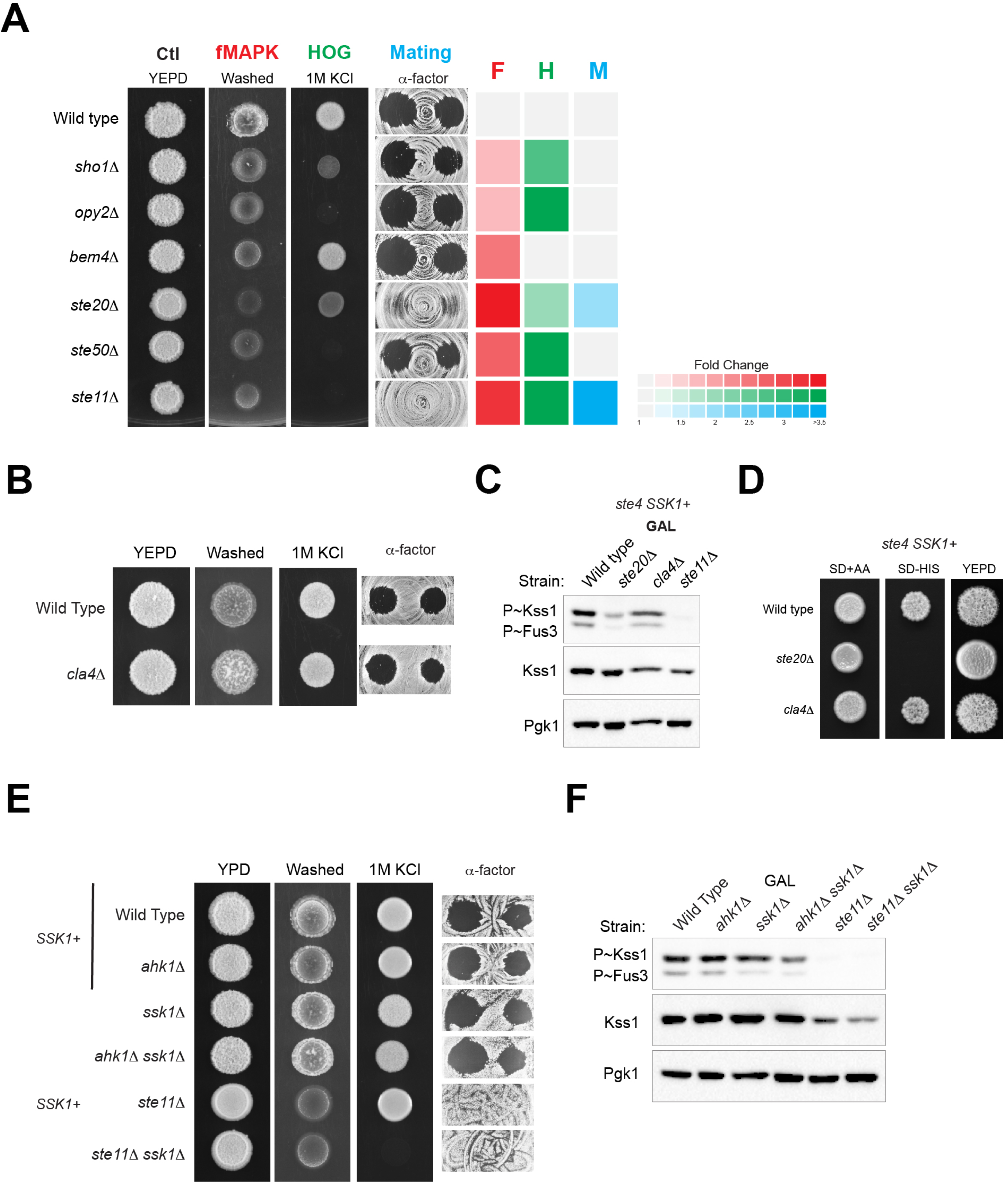
Role of different shared proteins in MAP kinase pathway activity. **A)** Phenotypes of different shared proteins mutants in tests for MAP kinase pathway activity. Color represents a phenotype in the PWA (fMAPK, F, red), growth on 1M KCl (HOG, H, green) or halo formation in response to α-factor (Mating, M, blue). Intensity represents the severity of the phenotype based on quantitation by ImageJ analysis; grey, no phenotypic difference from wild type. **B)** Phenotypes of wild-type cells and *cla4*Δ mutant by tests described in panel Fig. S3A. **C)** Immunoblot analysis of the *cla4*Δ mutant and controls. **D)** Comparison of wild-type cells to the *ste20*Δ and *cla4*Δ mutants by the *FUS1-HIS3* growth reporter and colony morphology after 4 d growth on YEPD. **E)** Phenotypes of the indicated combinations of the *ahk1*Δ mutant and controls by tests described in panel Fig. S3A. **F)** Immunoblot analysis of the *ahk1*Δ mutant and controls using the indicated antibodies.

**Figure S4.**
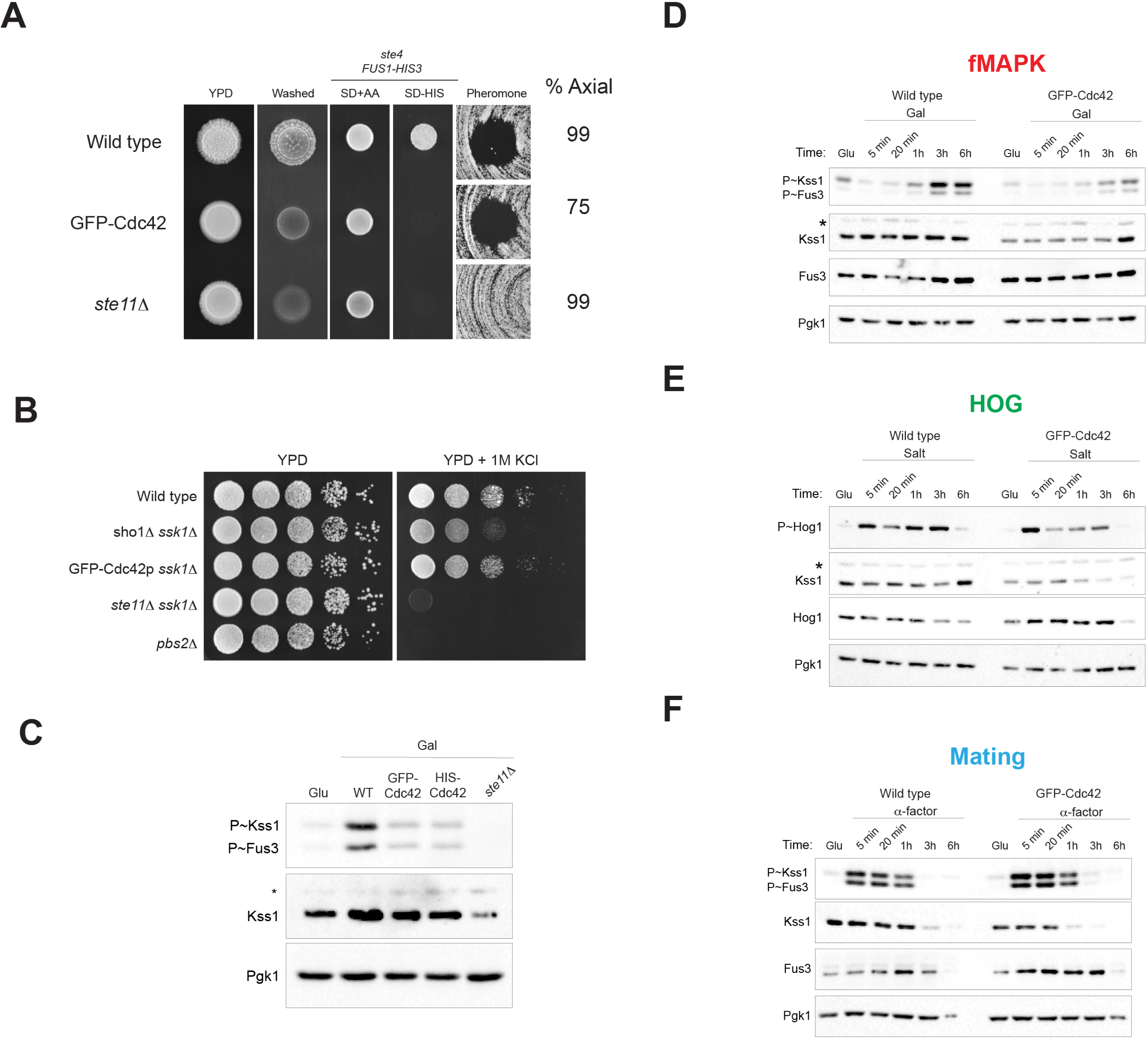
Functional analysis of N-terminally tagged versions of Cdc42p. **A)** Wild-type and *cdc42::NAT* cells carrying pGFP-Cdc42p or the *ste11*Δ mutant were spotted onto YEPD and examined by the PWA. Cells were spotted onto SD+AA (control) and SD–HIS media to assess the *FUS1-HIS3* reporter (strains are *ste4*Δ). Halo assays were performed on YEPD + 1.2µM of α-factor. Numbers indicate percentage of cells budding axially, determined by DIC microscopic examination of cells grown to mid-log phase. **B)** Serial dilutions of wild-type cells or cells with indicated gene knockouts were spotted onto YEPD or YEPD + 1M KCl for 2 d. **C)** Immunoblot assays of phosphorylated (P∼) MAP kinases in cell carrying pHis-Cdc42p or pGFP-Cdc42p with genomic *cdc42::NAT* (PC6685), and *ste11::NAT* (PC 6605). Cells were grown to mid-log phase in YEPD and harvested as a control (Glu) or induced with YEP-Gal for 3 h and harvested (Gal) for protein extraction. The asterisk indicates a background band seen in some blots. **D)** Immunoblot analysis measuring P∼Kss1p and P∼Fus3p levels in wild type cells and cells carrying pGFP-Cdc42p. Cells were grown to mid-log phase in YEP-GAL to induce fMAPK pathway. **E)** Immunoblot analysis of P∼Hog1p in wild type cells and cells carrying pGFP-Cdc42p induced with 0.4M KCl. **F)** P∼Kss1p and P∼Fus3p levels in wild type cells and cells carrying pGFP-Cdc42p induced with 1.2µM of α-factor. In the three cases, immunoblots were also probed with antibodies specific for Fus3p, Kss1p, and Hog1p. Anti-Pgk1p antibodies were used as a control for protein levels. Asterisk, background band.

**Figure S5.**
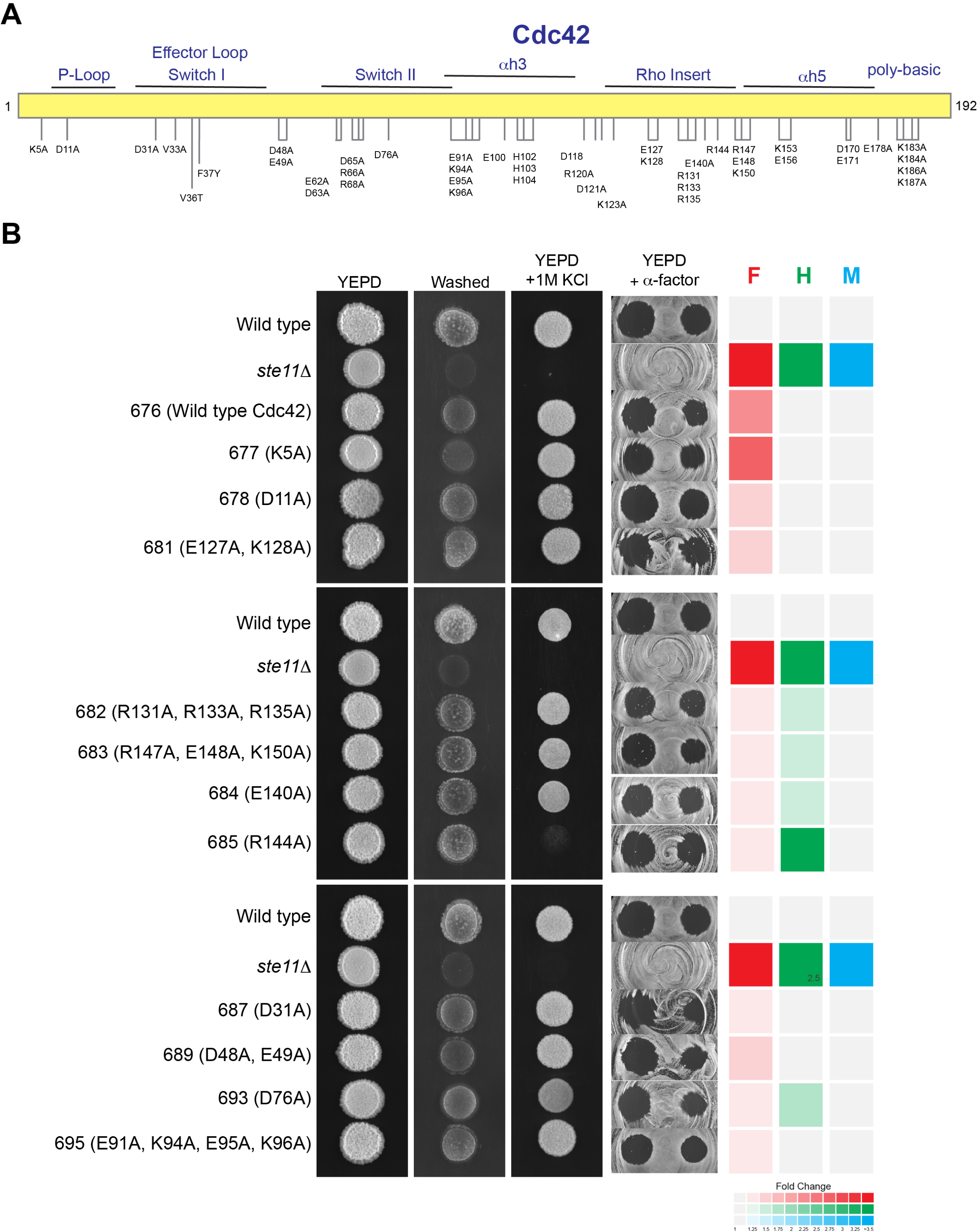
Analysis of *CDC42* alleles by functional assays of Cdc42p-dependent MAPK pathways. **A)** Diagram of Cdc42p with alleles tested in the study, the scheme represents the different parts of Cdc42p. **B)** Equal amounts of wild-type cells (PC6810), *ste11*Δ (PC6605) and cells carrying the indicated alleles of *cdc42* were spotted onto YEPD, YEP-Gal, or YEPD + 1M KCl media. Plates were incubated for 2 d and photographed. To evaluate invasive growth PWA experiment was performed. For halo assays, cells were spread on YEPD media followed by addition of 3 µl and 10 µl of 1mg/ml α-factor. Color intensity represents output phenotypes for MAPK pathways by ImageJ analysis; grey, no effect.

**Figure S6.**
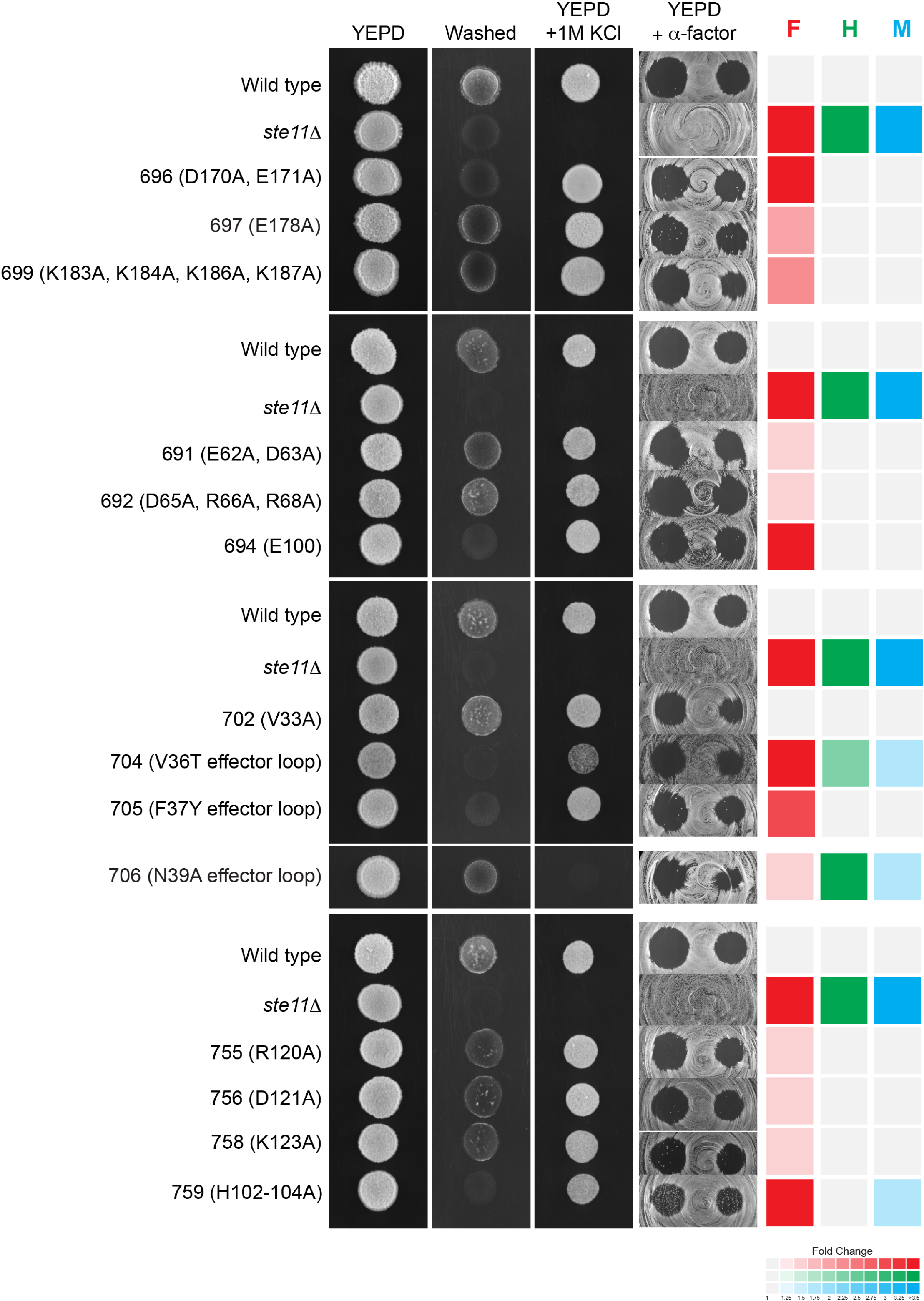
Analysis of *CDC42* alleles by functional assays of Cdc42p-dependent MAPK pathways. Cells were tested as indicated in Figure S5.

**Figure S7.**
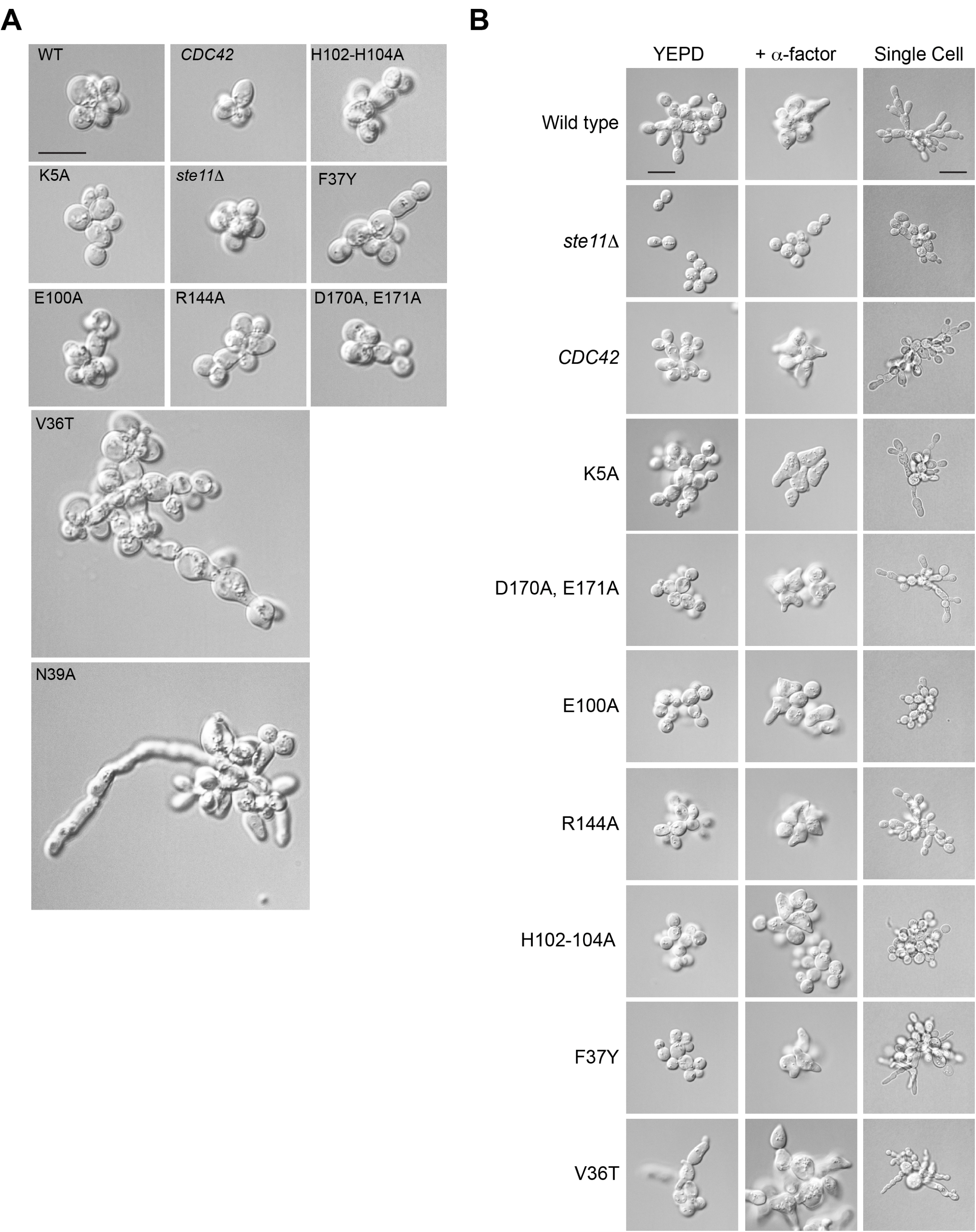
Cell morphology of *CDC42* alleles. **A)** Wild-type and cells carrying the indicated *CDC42* allele were grown to mid-log phase and photographed. Cells were examined by DIC microscopy at 100X. Bar, 10 microns **B)** Shmoo formation and single cell assay. Wild-type cells (PC6810), the *ste11*Δ (PC6605) mutant and cells carrying indicated alleles were grown to mid-log phase in YEPD, washed with water and resuspended in YEPD (control) or YEPD with 6 µM α-factor for three hours to examine shmoo formation. Cells were examined by DIC microscopy at 100X. Bar, 10 microns. *cdc42-130*^F37Y^ had ∼5% of cells with an irregular morphology in YEPD (*Table S1*). For single cell assay (last panel) 1 ml of saturated culture was washed and resuspended in water. Three microliter of water suspension was diluted in 400 µl water and spread on medium lacking glucose. A drop of 20 µl of 20% galactose was added off-center edge to the plate. Bar, 10 microns.

**Figure S8.**
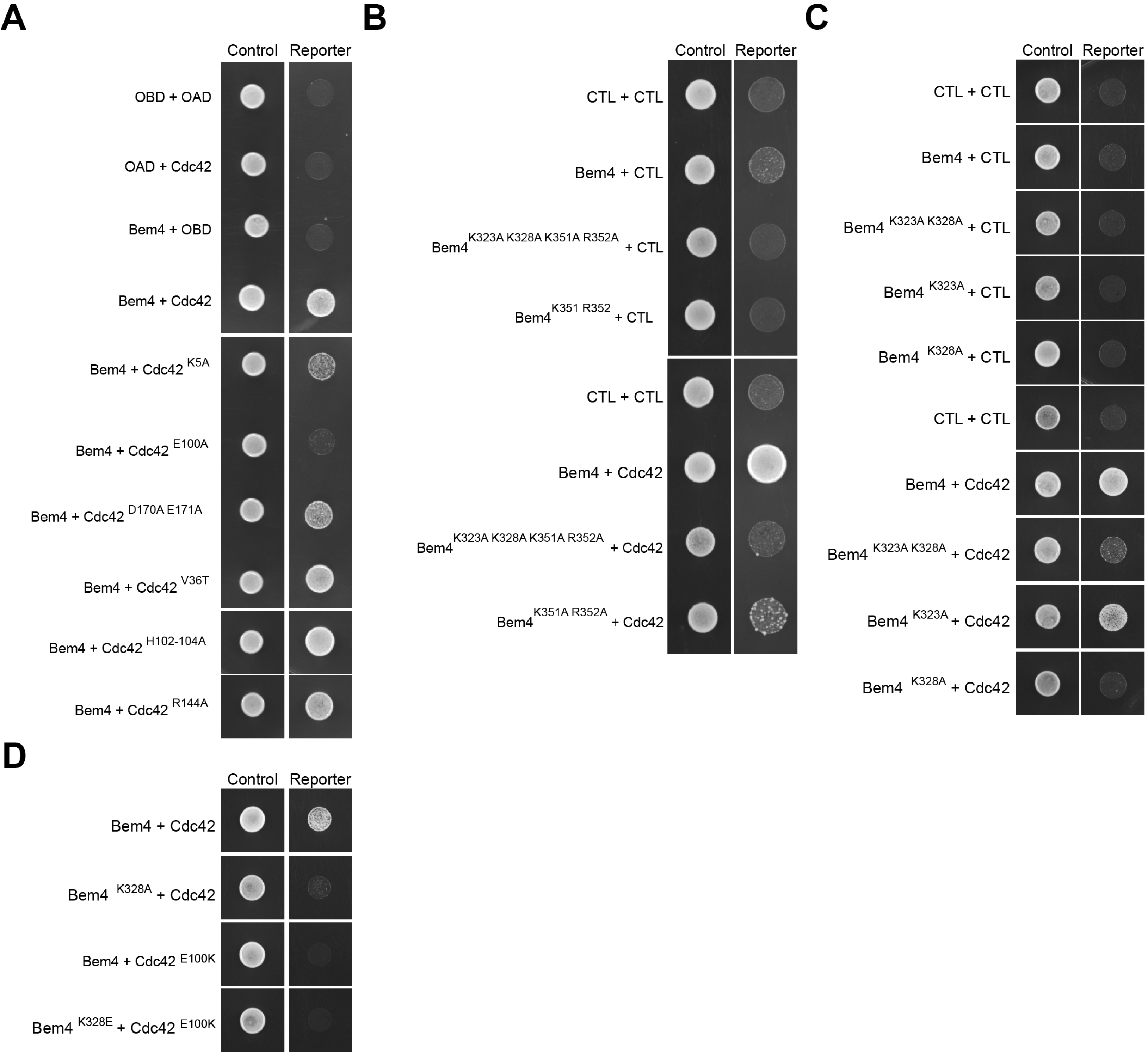
Interaction between Bem4p and versions of Cdc42p. **A-D)** Two-hybrid analysis of the indicated versions of Cdc42p and Bem4p. Cells carrying the indicated plasmids were tested in control (SD) and reporter (SD-HIS) plates; plates were photographed after 2 d.

**Figure S9.**
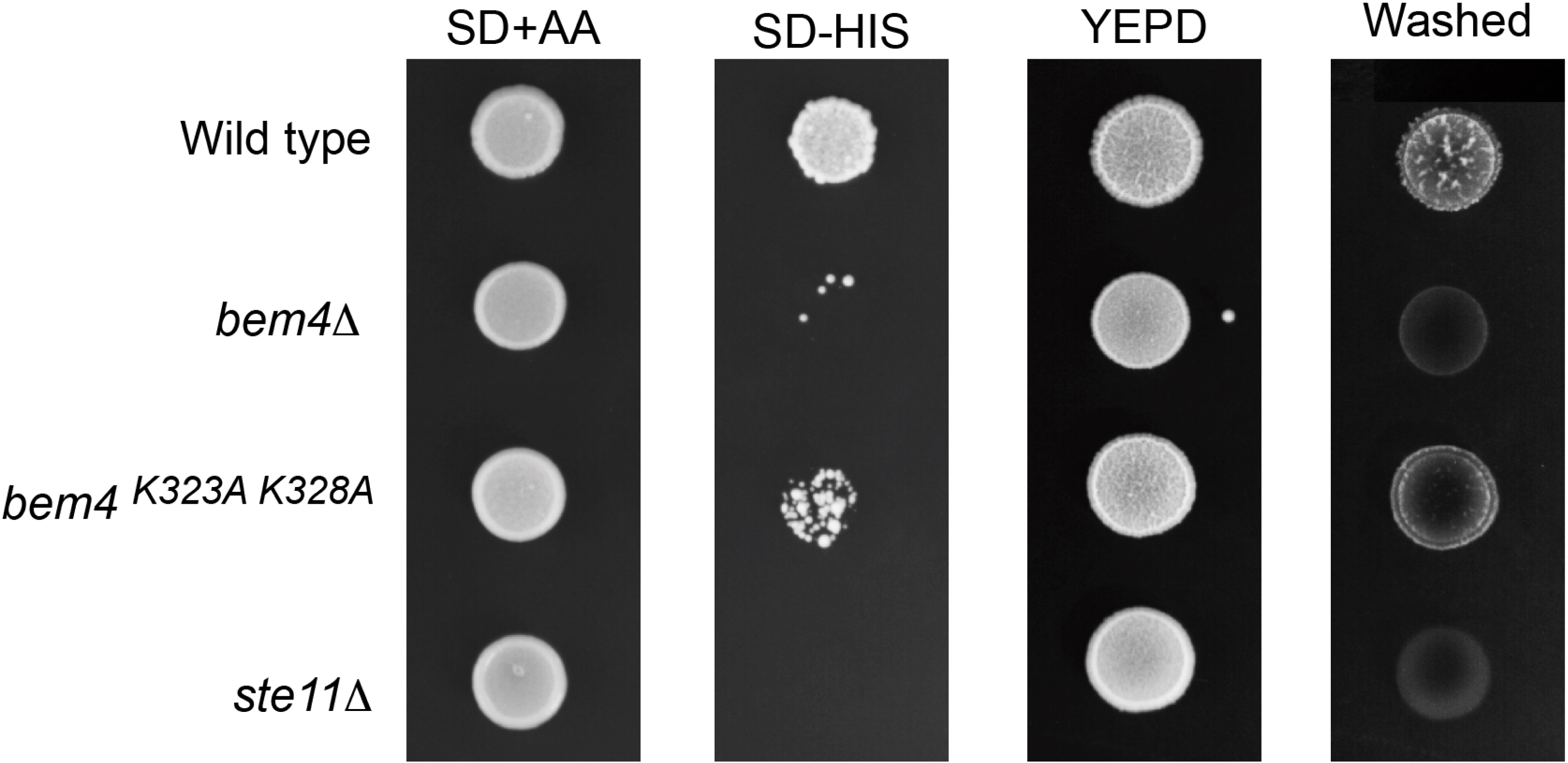
Role of Bem4^K323A,328A^ version in regulating fMAPK pathway. The two-point mutations of Bem4p ^K323A K328A^ were inserted in the genomic DNA of the wild-type (PC538). Cells were spotted on SD (control) and SD-HIS (reporter) to evaluate the activity of the *FUS1-HIS* reporter. For the PWA, cells were grown in YPED media for 48 h.

**Figure S10.**
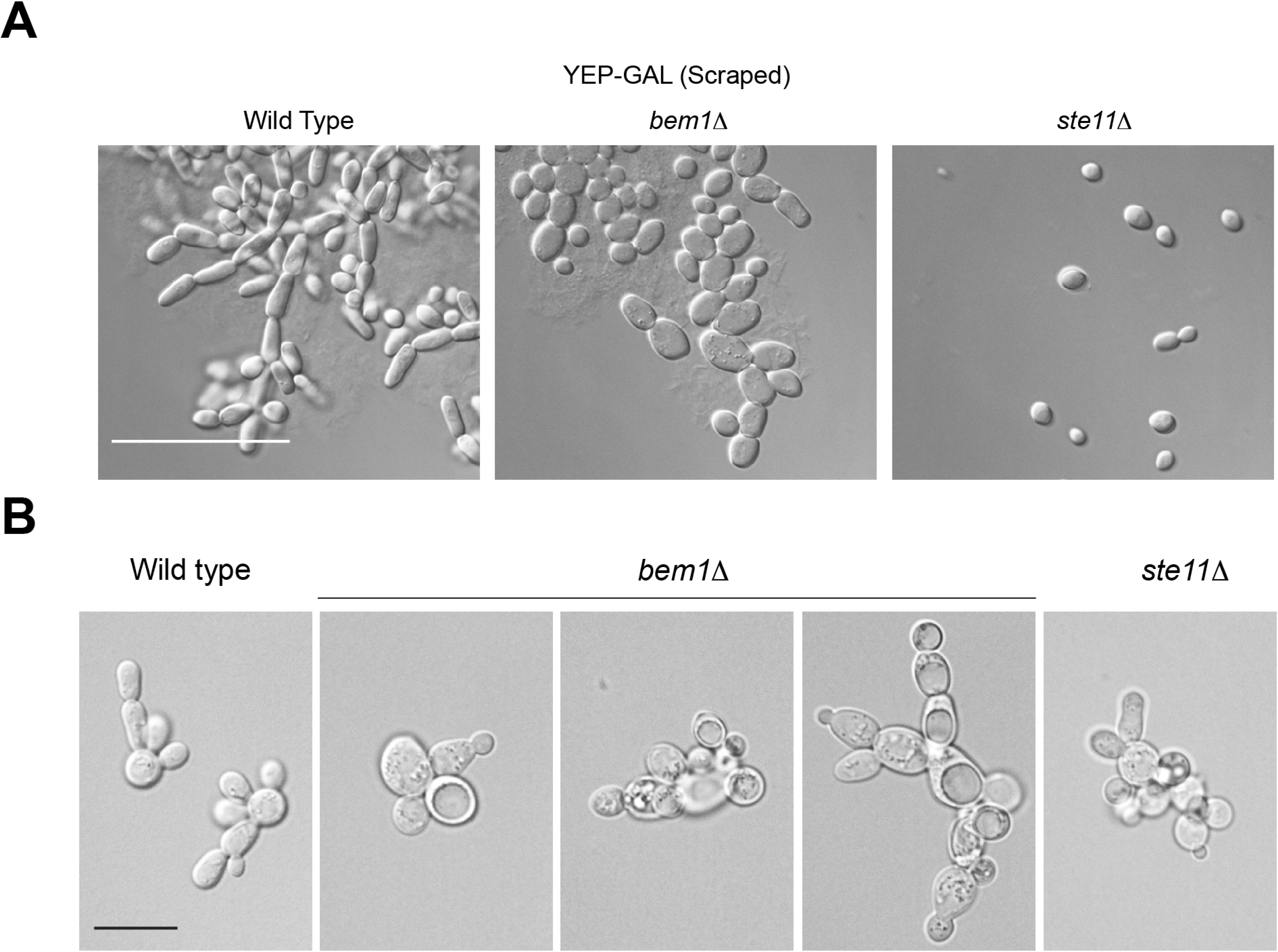
Morphology of *bem1* mutant and controls under fMAPK pathway inducing conditions. **A)** Wild-type cells (PC6603), *bem1*Δ and *ste11*Δ mutants were grown in YEP-GAL plates for 48h and then observed by microscopy. **B)** Indicated strains were grown to mid log phase and examined by DIC microscopy at 100X. Bar, 10 microns.

**Figure S11.**
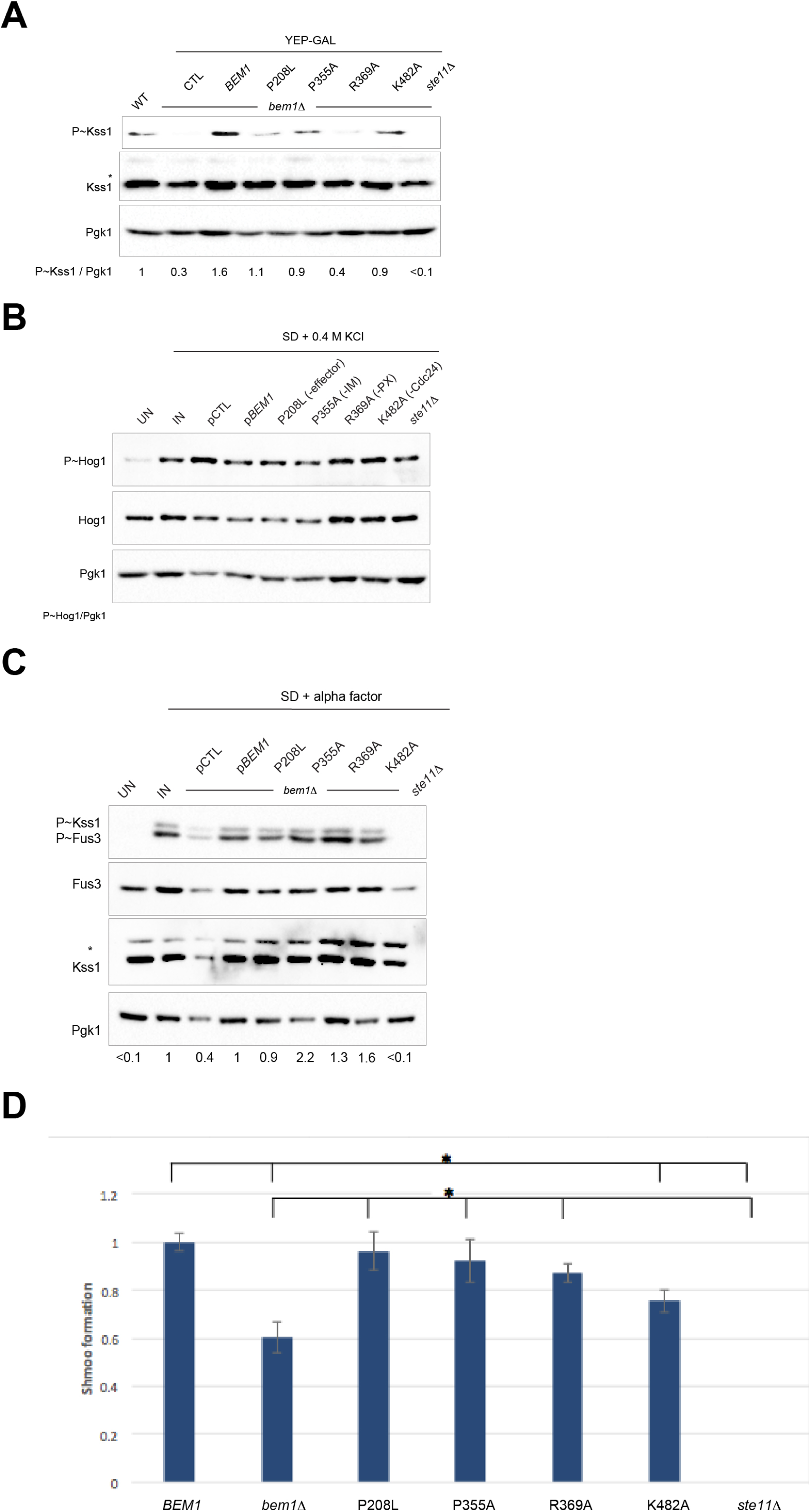
Role of *BEM1* alleles in regulating Cdc42p-dependent MAPK pathways. **A)** Immunoblot analysis of P∼Kss1p levels in response to galactose in wild-type cells, *bem1*Δ mutants carrying the indicated *BEM1* allele, and *ste11*Δ mutant. **B)** P∼Hog1p levels in response to 0.4M KCl in the same cells explored in panel S11A. **C)** Examining P∼Fus3p and P∼Kss1p levels in response to α-factor induction in the same cells explored in panel S11A In the three cases, specific antibodies were also used to detect MAP kinase levels and Pgk1p. Numbers indicate relative band intensity of P∼MAPK to total MAPK levels normalized to the control condition, which was set to 1. **D)** Relative quantification of shmoo formation in the indicated strains.

**Figure S12.**
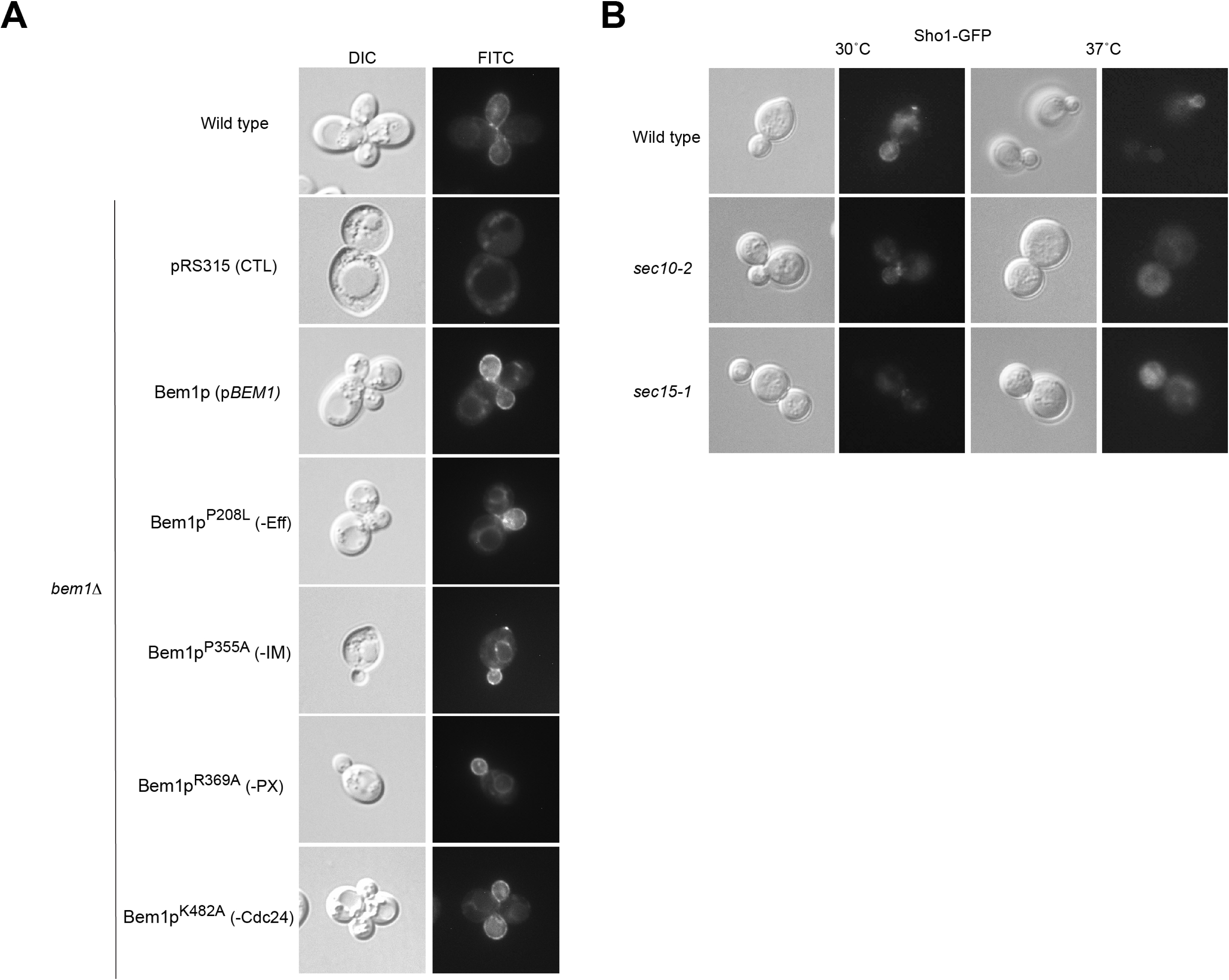
Role of *BEM1* alleles in Sho1p-GFP localization. **A)** Sho1p-GFP localization in wild-type cells (PC6603) and the *BEM1* alleles strains. Cells were grown in YEPD media and examined by DIC and fluorescence microscopy at 100X. **B)** Sho1p-GFP localization in wild-type cells (PC6603) and cells carrying indicated temperature sensitive alleles of secretory pathway mutants (PC1660, PC1661). Cells grown in YEPD at 30°C or 37°C as indicated. Bar, 10 microns.

**Figure S13.**
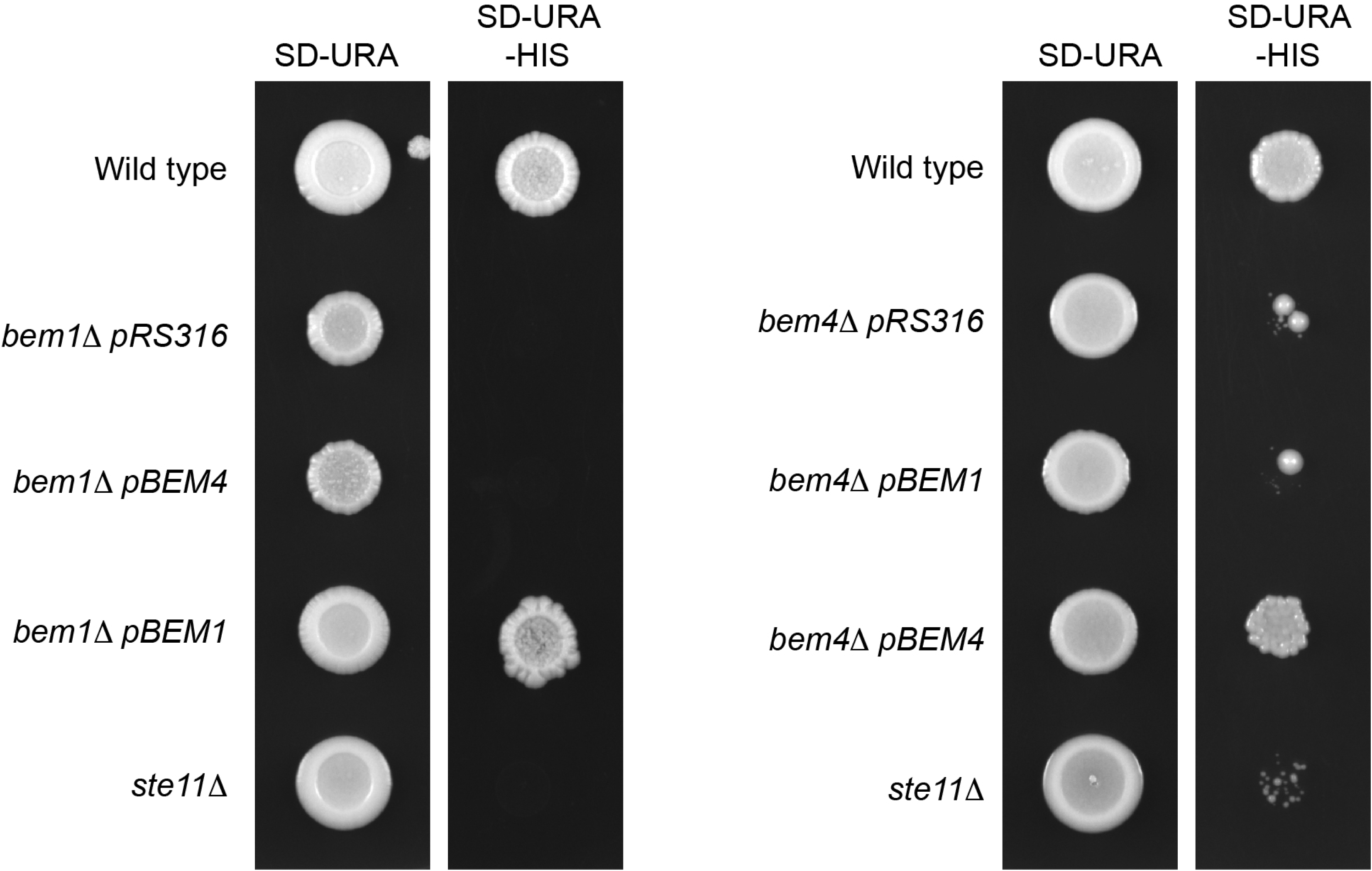
Comparison of the signaling defect of the *bem1*Δ and *bem4*Δ mutants. Overexpression of *BEM4* and *BEM1* in the *bem1*Δ mutant to examine signaling redundancy. Wild type (PC538), *bem1*Δ (PC6509) carrying indicated plasmids, and the *ste11*Δ mutant (PC3861) were spotted on SD-URA, SD-URA-HIS to examine *FUS1-HIS3* reporter activity. At right, the same experiment was performed in the *bem4*Δ mutant.

***Movie S1*.** Modeling the interaction between Bem4p and Cdc42p.

**Table S1.**
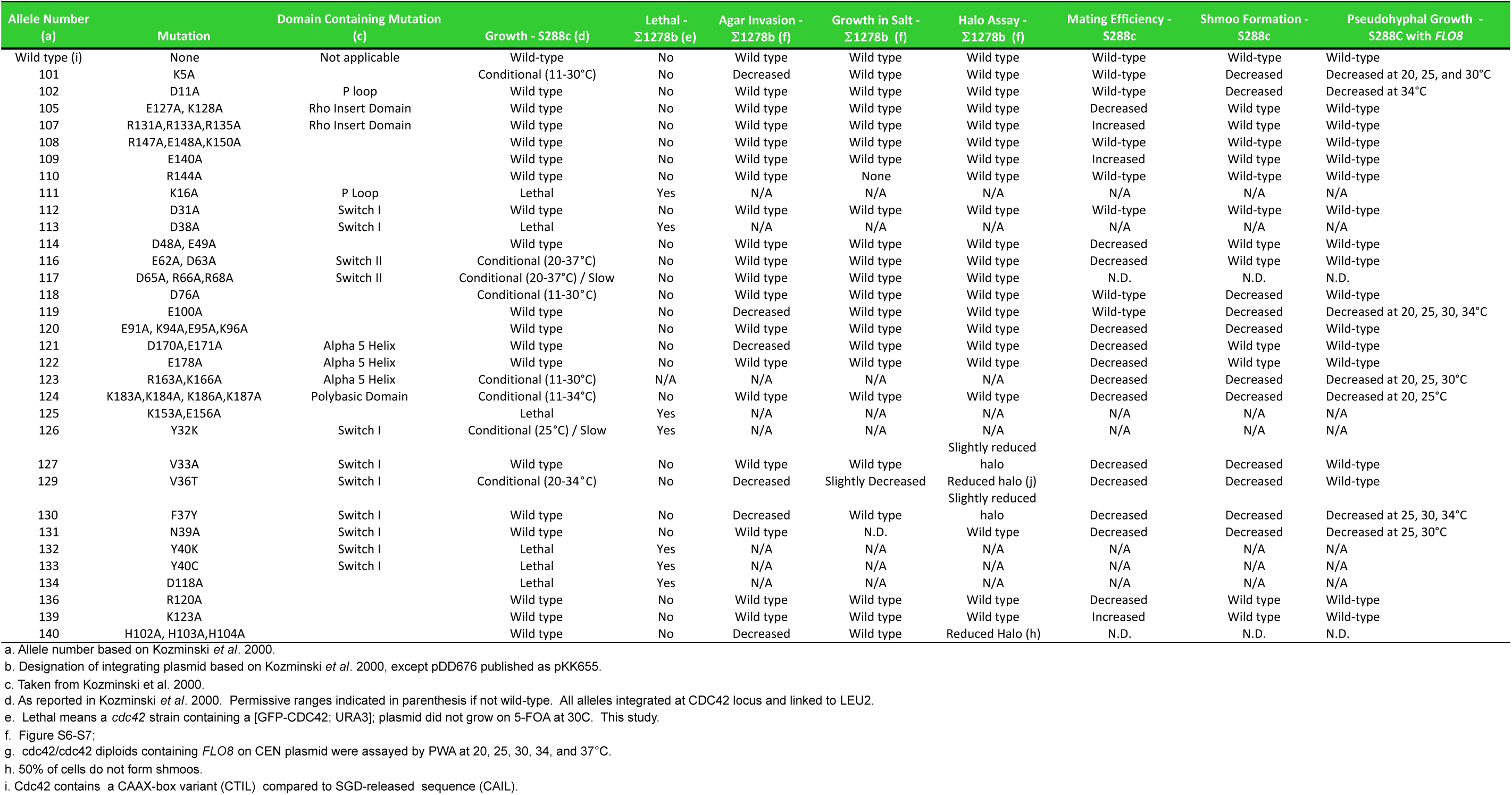
Analysis of the roles of *cdc42* alleles in regulating MAPK pathways

